# Improving efficiency of homology-directed repair with ZIP CRISPR

**DOI:** 10.1101/2025.03.13.642982

**Authors:** C Thibault, J Rosier, R Espasa, V Marin, M Riandière, C Berges, S Fayet, I Lamrissi-Garcia, M Lalanne, T Trian, S Bui, S Dabernat, J Boutin, F Moreau-Gaudry, A Bedel

## Abstract

Precise genome editing technologies create the potential for genetic studies and innovative gene therapies. Here we present new CRISPR-Cas9 tools, named ZIP CRISPR, loaded with a single-stranded oligodeoxynucleotide (ssODN) template on the Cas ribonucleoprotein complex. The ssODN template is annealed to an extended guide RNA (gRNA) allowing its nuclear delivery at the right place, i.e. the targeted DNA cut, and at the right time. This new template import system is easy-to-design, easy-to-use, inexpensive and versatile. It increases homology-directed repair (HDR) editing efficiency using Cas9 nuclease up to 12-fold with a mean increase of 5-fold as demonstrated at many *loci* in many cell types. Based on a heteroduplex gRNA-ssODN, it can also be used with the Cas9 nickase, resulting in HDR editing with minimal InDels, and preventing double-strand break (DSB)-mediated genotoxicity. ZIP CRISPR is a non-viral platform adaptable to targeted DSB (higher HDR editing efficiency) or nick (higher safety) to precisely model and correct a wide range of edits. It is suitable for many biological applications and could be considered for HDR-based gene therapies.

## Introduction

The CRISPR-Cas9 system induces DNA double-strand breaks (DSBs) at targeted sites to activate mainly two competitive DNA repair pathways: homology-directed repair (HDR), which allows precise editing with a homologous DNA template, and non-conservative NHEJ/MMEJ pathways, which connect two ends of the broken DNA and are often accompanied by random insertions and deletions (InDels)^1^. The relatively low efficiency of HDR compared to NHEJ/MMEJ repair has been a major hurdle in achieving precise genome editing (PGE) at desired *loci*. Because of their higher efficacy, clinical genome editing protocols have been restricted to imperfect but efficient NHEJ-based approaches. Several techniques have been proposed to obtain a higher PGE rate, in particular by modulating repair pathways, either by pharmacological NHEJ/MMEJ inhibition^2–6^, or by HDR activation (*via* the expression of HDR actors^7–9^, Cas9-fusion proteins to enhance the HDR pathway at the targeted *locus*^10–13^ or S/G2 synchronization^14^). While promising, these approaches may alter the normal behavior of the targeted cells, raising substantial safety concerns when explored in a therapeutic context^15^. In addition, HDR remains to be optimized for therapeutic applications. The presence of the exogenous DNA template at the DNA cut is critical for HDR editing. For example, fluorescently labeled donor DNA can be used to enrich cells that are likely to be edited *via* HDR^16,17^. Several teams have already done important work to develop Cas9 tools to directly carry the single-stranded oligodeoxynucleotide (ssODN) template on the RNP complex. ssODN donors can be tethered to the Cas9 protein *via* the Cas9-avidin biotin-ssODN system^11^, the RNP-DNA (RNPD) system^18^, the Cas9-PCV system^19^, the Cas9-AeF DBCO-adaptor ssODN system^20^, or a platform that relies on thiol-maleimide chemistry and DNA-base pairing^21^. Alternatively, ssODN can be tethered to the gRNA *via* a S1mplex system and a biotinylated ssODN^22^, either with a chemical link between tracrRNA and the ssODN^17^ or directly with a chimeric RNA-DNA donor guide^23^. However, they mostly rely on chemical modifications of the Cas9, gRNA, ssDNA donor and/or used fused protein. Thus, their uses are constrained by chemical synthesis difficulties and risk of destabilizing the Cas9-gRNA interaction. We therefore sought to develop a chemical modification-free approach that can recruit ssODN donors to the target site and enhance HDR efficiency. Here, we describe the development of new efficient CRISPR-Cas9 tools, named ZIP CRISPR, that carry a ssODN template annealed to a modified gRNA allowing ribonucleoprotein (RNP) stability. This specific gRNA-ssODN heteroduplex is easy-to-design and stable within cells after transfection. It does not require any specialized and/or expensive chemical modifications or modified Cas9. The simplicity of using unmodified Cas9, ssODN and easily available modified gRNA makes it readily transferable to all laboratories. Importantly, it is versatile, allowing increased HDR editing efficiency at many *loci*, in cell lines and primary cells with nuclease. Interestingly, this strategy also unlocked HDR editing by using safer single nickase.

## Results

### ZIP CRISPR improves precise genome editing (PGE) using Cas9 nuclease

To increase the effective concentration of the donor template at the DSB and thus HDR editing efficiency, we developed a strategy to hybridize the ssODN donor template to the gRNA. For that, we added a 3’-extension (light pink in the schema) on the gRNA. This 20nt extension has the same sequence as the 3’-extension of the first versions of pegRNAs published by Anzalone *et al.* in 2019^24^, as we hypothesized that it will not destabilize the gRNA structure. This extension is complementary to a 5’-extension (light blue in the schema) of the 80nt ssODN template for annealing (Fig. 1a). The RNP complex formed is named ZIP and the control RNP with regular free ssODN and gRNA is named UNZIP. To quantify CRISPR-Cas9-mediated genome editing, we used a rapid flow cytometry assay based on eGFP to BFP conversion^25^ (Fig. 1b/c). In the case of HDR editing, the substitution of a histidine in place of a tyrosine at position 66 in the chromophore of eGFP shifts its emission toward the blue spectrum^26^ (Fig. 1b). HEK293T cells, K562 cells and human foreskin fibroblasts (hFFs) were stably transduced at low multiplicity of infection with a lentiviral construct containing an eGFP/puromycin resistance cassette to obtain less than 10% of eGFP^+^ cells. After selection, we obtained cells with one copy of eGFP per cell. Reliable simultaneous quantification of HDR– and imprecise-editing was obtained by flow cytometry one week after editing by measuring the appearance of BFP fluorescence or the loss of eGFP fluorescence, respectively (Fig. 1c). The BFP detection is highly specific to HDR editing (no BFP without template). The eGFP fluorescence loss is mainly due to imprecise editing but can rarely be related to a spontaneous loss of fluorescence or to untransduced cells. Interestingly, overall editing efficiency was not affected by the extended gRNA but PGE (HDR editing) efficiency was significantly increased by 2-fold with ZIP compared to UNZIP nucleofection in eGFP^+^-HEK293T cells (14.9% ± 0.7 *vs* 8% ± 1 respectively, Fig. 1d and Supplementary Fig. S1a) and was confirmed in an eGFP^+^-K562 cell line (10.2% ± 3.8 *vs* 5.2% ± 2.6 respectively, Fig. 1e). These data obtained in two cell lines demonstrated that ZIP CRISPR increases PGE by using CRISPR-Cas9 nuclease, with a 2-fold increase in the HDR/InDels ratio. To determine if this increase was really due to an improved proximity between the ssODN template and the gRNA, we performed a FRET experiment using a Cy3-ssODN (containing the hybridization sequence) and a Cy5-probe annealed to the gRNA (containing or not the hybridization sequence) (Fig. 1f). We analyzed FRET signals (Cy5 emission after Cy3 excitation) of RNP complex either directly by spectrofluorometry or after cell transfection by confocal microscopy (Supplementary Fig. S1b). Using spectrofluorometry, we observed an increase by 2.4-fold in FRET signal (p= 0.1) with the ZIP compared to UNZIP condition (Supplementary Fig. S1c). This means that the proximity between ssODN and gRNA seems to be increased with the ZIP system, in favor of hybridization. We confirmed this amplification of the FRET signal by confocal microscopy, seven hours after cell transfection with the ZIP compared to the UNZIP system. Indeed, the number of FRET-positive pixels for 100 cells is increased when gRNA and ssODN are hybridized using the same doses as for editing experiments (Fig. 1g/h left panel). These data confirm ZIP RNP stability into cells after 37°C culture. To test the stability of the gRNA-ssODN annealing at low concentration, we used a lower dose (⅓ of the usual dose) and confirmed a rise in the number of FRET-positive pixels for 100 cells under the ZIP condition (Fig. 1h right panel). Surprisingly, by confocal microscopy analysis, we observed at usual and low doses, while using the same amount of components, that the Cy signals are higher in the ZIP condition (Fig. 1g and Supplementary Fig. S1d). This could be due to a better transfection efficiency or an increased stability when gRNA and ssODN are annealed. However, when normalizing on Cy3/Cy5 transfection efficiency, FRET-positive pixels are still increased in the ZIP condition (Supplementary Fig. S1e). Taken together, these results confirmed that gRNA and ssODN remain attached or at least sufficiently closed in the cells at 37°C seven hours after transfection, thereby supporting PGE in the ZIP condition.

**Fig. 1:**
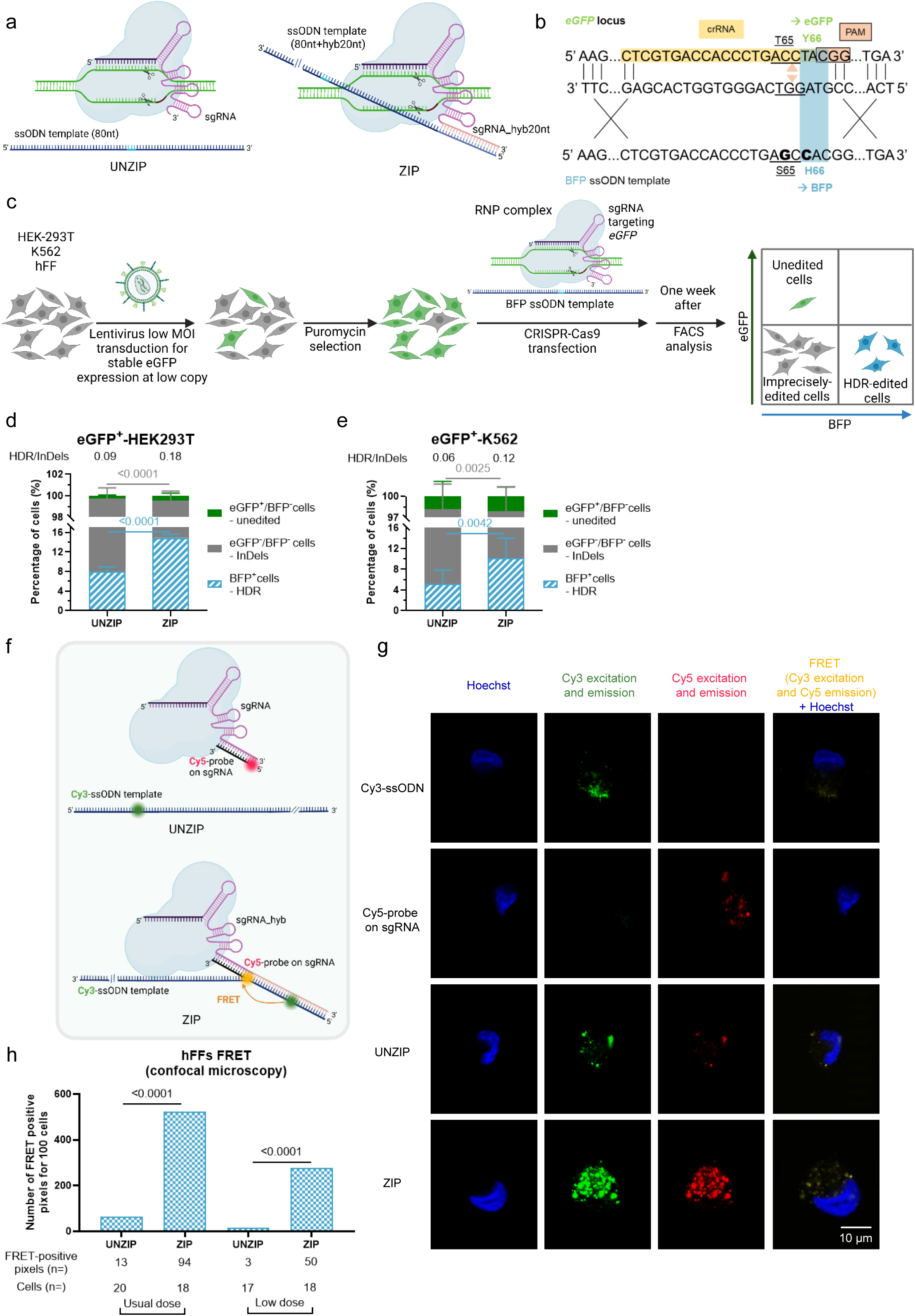
Set up and validation of ZIP CRISPR to increase HDR editing efficiency with Cas9 nuclease in cell lines. **a** Representation of RNPs UNZIP (Cas9 nuclease with free ssODN template) and ZIP (Cas9 nuclease with ssODN template import: elongated ssODN – in light blue-for hybridization to the elongated gRNA – in light pink-). **b** Editing strategy to convert eGFP into BFP. T65S and Y66H amino acid substitutions correspond to a shift in the fluorescence excitation and emission spectra of the protein, converting eGFP to BFP. **c** Construction of cell models and result analysis timeline. **d and e** Flow cytometry quantification of the edition with Cas9 nuclease with (ZIP) or without (UNZIP) the correction template physically bound to the gRNA in eGFP^+^-HEK293T cells (n=4) and eGFP^+^-K562 cells (n=8), respectively. The proportions of cells expressing BFP (HDR-edited), eGFP (unedited) or neither of these fluorophores (imprecisely edited, with InDels) one week after transfection are reported. The mean ± SD is shown. The HDR/InDels ratio is indicated at the top of the graphs. Statistical significance determined by two-way ANOVA. **f** Representation of RNPs UNZIP and ZIP used for the FRET experiments. **g** Illustrative confocal microscopy imaging of hFFs transfected with either Cy3-ssODN, Cy5-probe on sgRNA, UNZIP or ZIP systems at usual dose. Hoechst, Cy3, Cy5 and FRET merged with Hoechst signals are illustrated for each condition. **h** Quantification of FRET signals observed by confocal microscopy in hFFs transfected with ZIP or UNZIP systems at usual or low dose (⅓ less) (n=1). The numbers of FRET-positive pixels normalized for 100 cells are represented. The exact number of FRET-positive pixels and cells analyzed are indicated at the bottom of the graph. Statistical significance determined by Chi-square with Yates’ correction. Schemas created with BioRender.com.

### ZIP CRISPR’ improvements

To establish whether ssODN template stability and quantity were critical for PGE, we designed two new ZIP CRISPR systems with either double ssODN hybridization on gRNA or elongated ssODN (120nt) to slow down its degradation. We named these new systems ZIP_Double_Lock and ZIP_Long_Lock (Supplementary Fig. S2a). Indeed, because these modifications could affect gRNA stability and Cas9 interaction, we added a t-lock element (orange in the schema) in the second loop of the gRNA, as previously described^27^ to reinforce its stability. This t-lock element alone (control ZIP_Lock, Supplementary Fig. S2a) did not modify editing efficiency but ZIP_Double_Lock, and more efficiently ZIP_Long_Lock improved PGE in eGFP^+^-K562 cell line with respectively +37.2% and +70% of HDR-edited cells (Fig. 2a and Supplementary Fig. S2b/c). For the ZIP_Double_Lock, the lower efficiency compared to the ZIP_Long_Lock could be due to a steric hindrance when two ssODN are present in the RNP complex. The important effect with ZIP_Long_Lock was due to elongation (control ZIP_Long, +47.3%) combined with the t-lock loop. To create these new systems, we elongated the extension length to 30nt. To determine if the HDR increase observed could have been due to a change in the extension/annealing length (from 20 to 30nt for ZIP_Long and ZIP_Long_Lock and from 20 to 2*15nt for ZIP_Double_Lock), we created new versions of ZIP_Long_Lock with annealing of 15nt, 20nt or 30nt and demonstrated that this change in annealing length doesn’t impact HDR editing efficiency (Supplementary Fig. S2d/e), suggesting that the increase observed with ZIP_Long_Lock compared to ZIP is due to the t-lock element in the gRNA and the elongation of the ssODN template but not to the 30nt for annealing. Because template length seems to be an important parameter, we again elongated the ssODN template (170nt, ZIP_ExtraLong_Lock, Supplementary Fig. S2f). We also tested the putative impact of hybridization position in the template. For that, we flipped the template with a 3’-extension for hybridization to the gRNA (ZIP_Inv, Supplementary Fig. S2f). However, neither of these constructs improved HDR editing efficiency compared to ZIP_Long_Lock (Supplementary Fig. S2g/h/i). Altogether, ssODN elongation (120nt) associated with a t-lock loop in the gRNA (ZIP_Long_Lock) was the most efficient system, so we named it ZIPmax and used it for the following experiments. To check if the increase in HDR editing with ZIPmax was due to the initial extension sequence chosen, we randomly created a new sequence with the same nucleotides and compared this ZIPmax_V2 to ZIPmax. We did not observe any difference in terms of HDR-editing suggesting that the sgRNA-ssODN hybridization, no matter the sequence, is important in ZIP CRISPR (Supplementary Fig. S2j/k). We decided to keep the initial sequence for the following experiments.

**Fig. 2:**
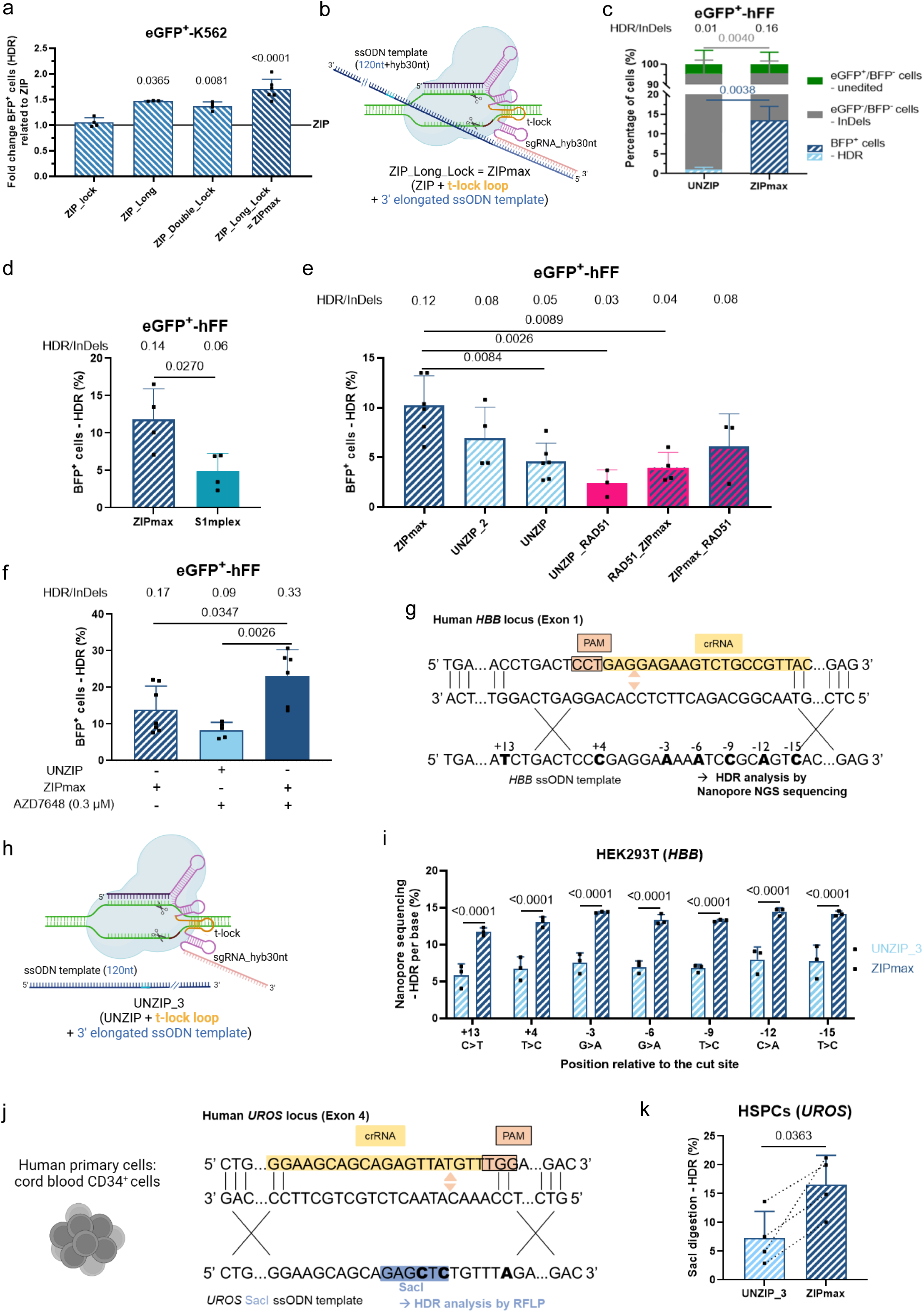
Improvement and benchmarking of ZIP CRISPR to boost HDR editing with Cas9 nuclease in cell lines and primary cells. **a** Flow cytometry quantification of the HDR edition with Cas9 nuclease with ZIP_Lock (n=4), ZIP_Long (n=3), ZIP_Double_Lock (n=4) and ZIP_Long_Lock (n=6) in eGFP^+^-K562 cells. The proportions of cells expressing BFP one week after transfection are reported. Fold-changes indicate the fold-increase between the ZIP_Lock, ZIP_Long, ZIP_Double_Lock and ZIP_Long_Lock systems relative to ZIP. The mean ± SD is shown. Statistical significance to ZIP determined by Kruskal-Wallis test. **b** Representation of ZIP_Long_Lock=ZIPmax system. **c** Flow cytometry quantification of the edition with Cas9 nuclease with ZIPmax or UNZIP in eGFP^+^-hFFs (n=5). The proportions of cells expressing BFP, eGFP or neither of these fluorophores one week after transfection are reported. The mean ± SD is shown. The HDR/InDels ratio is indicated at the top of the graphs. Statistical significance is determined by two-way ANOVA. **d** Flow cytometry quantification of the edition with Cas9 nuclease with ZIPmax or S1mplex in eGFP^+^-hFFs (n=4). The proportions of cells expressing BFP one week after transfection are reported. The mean ± SD is shown. The HDR/InDels ratio is indicated at the top of the graphs. Statistical significance is determined by unpaired t-test. **e** Flow cytometry quantification of the edition with Cas9 nuclease with ZIPmax (n=6), UNZIP_2 (n=4), UNZIP (n=6), UNZIP_RAD51 (n=3), RAD51_ZIPmax (n=4) or ZIPmax_RAD51 (n=3) in eGFP^+^-hFFs. The proportions of cells expressing BFP one week after transfection are reported. The mean ± SD is shown. The HDR/InDels ratio is indicated at the top of the graphs. Statistical significance is determined by one-way ANOVA. **f** Flow cytometry quantification of the HDR edition with Cas9 nuclease with UNZIP or ZIPmax and with or without inhibition of NHEJ repair with AZD7648 in eGFP^+^-hFFs (n=6). The proportions of cells expressing BFP one week after transfection are reported. The mean ± SD is shown. The HDR/InDels ratio is indicated at the top of the graphs. Statistical significance determined by one-way ANOVA. **g** Editing strategy to edit *HBB* in HEK293T cells. **h** Representation of the UNZIP_3 system. **i** Nanopore sequencing to quantify HDR editing per base with UNZIP_3 or ZIPmax (n=3). The mean ± SD is shown. Statistical significance determined by two-way ANOVA. **j** Editing strategy to edit *UROS* in CD34^+^ HSPCs. **k** Capillary electrophoresis of *SacI* digestion products to quantify HDR editing with UNZIP_3 or ZIPmax (n=4). The mean ± SD is shown. Statistical significance determined by unpaired t-test. Schemas created with BioRender.com.

### ZIPmax benchmarking and validation in endogenous *loci*

Then, we evaluated ZIPmax (Fig. 2b) in primary eGFP^+^-hFFs, which are difficult to edit. ZIPmax outperformed the UNZIP control by improving HDR editing efficiency up to 12-fold and the HDR/InDels ratio by 16-fold (Fig. 2c). Given this higher efficiency of ZIPmax, we decided to benchmark it to other strategies to improve HDR editing efficiency. First, we compared it to the S1mplex method proposed by Carlson-Stevermer *et al*. in 2017, because as ZIPmax it uses only commercially available components^22^. S1mplex is composed of a biotinylated ssODN and a gRNA modified to include an aptamer that binds streptavidin. We challenged our ZIPmax with S1mplex in eGFP^+^-hFFs and observed a 2.4-fold increase in HDR-editing using ZIPmax (Fig. 2d and Supplementary Fig. S3a). Then, we compared and combined ZIPmax to Jin *et al*.’s strategy that demonstrated that small ssODN extensions can recruit proteins and increase HDR editing. In particular, they demonstrated that a specific 24nt 5’-extension on the ssODN can recruit RAD51 and increase HDR in several cell types for several targets^7^. We compared ZIPmax with an UNZIP condition using an ssODN with or without this RAD51 recruitment motif, called UNZIP_RAD51 or UNZIP_2 respectively (Supplementary Fig. S3b). We observed that ZIPmax exceeds the UNZIP_RAD51 condition with a 4.1-fold increase (Fig. 2e and Supplementary Fig. S3c). Then, we tried to combine both approaches. As in ZIPmax the extension for annealing to the gRNA is also at the 5’ end, we created two versions with either RAD51 recruitment motif located in 5’ or 3’ position relative to the annealing extension. They are called RAD51_ZIPmax and ZIPmax_RAD51 respectively (Supplementary Fig. S3b). When combining RAD51 recruitment motif with ZIPmax, PGE remains lower than with the ZIPmax construction alone (Fig. 2e and Supplementary Fig. S3c).

Another way to increase the HDR/InDels ratio is to mitigate the DNA repair pathways. To date, the most potent molecule to inhibit the NHEJ pathway is AZD7648, a potent inhibitor of DNA-PKcs^3^. We decided to challenge ZIPmax with this compound. Using eGFP^+^-hFFs, ZIPmax provided a 1.7-fold higher precise editing efficacy compared to UNZIP with the optimal conditions of AZD7648 exposure (13.9% ± 6.5 *vs* 8.3% ± 12.1 respectively, Fig. 2f and Supplementary Fig. S4). Importantly, a complementary additive effect was observed with the simultaneous use of ZIPmax and AZD7648, leading to 23.1% ± 7.3% of HDR-edited cells. Given recent data showing the alarming genotoxicity of AZD7648^28,29^, we decided to use only ZIPmax without AZD7648 in the following experiments.

Next, we tested ZIPmax to edit the *HBB* endogenous *locus* (Fig. 2g) which is a pertinent genomic target for clinical applications. Because the t-lock element in the gRNA can improve HDR editing efficiency by itself at some *loci*^27^, we used the same gRNA (with t-lock stabilization and 3’-extension) as a control in UNZIP_3 (Fig. 2h). The only difference between UNZIP_3 and ZIPmax was the template, i.e. with or without the 5’-extension to anneal (or not) to the gRNA. NGS analysis by Nanopore sequencing across the targeted *HBB* region (−15 to +13nt around the cut site) showed that the seven nucleotides modified in the template were edited with a 2-fold increase in the HDR editing rate with ZIPmax (from around 7 to 13.5%, Fig. 2i).

To check the versatility of ZIP editing in clinically relevant cells, we evaluated it in cord blood-derived CD34^+^ hematopoietic stem and progenitor cells (HSPCs) by targeting the exon 4 of *UROS,* an enzyme of the heme biosynthesis (Fig. 2j). In these primary cells, PGE was measured by restriction fragment length polymorphism (RFLP). Digestion occurred after the perfect editing of two nucleotides to create a *SacI* restriction site. The HDR editing rate was more than 2-fold higher with ZIPmax than with UNZIP_3 (16.5% ± 5.1 *vs* 7.3% ± 4.6 respectively, Fig. 2k).

Taken together, these data showed the higher efficacy of ZIPmax in cell lines and primary cells to improve HDR editing efficiency for different targets.

### *CFTR* gene editing in human pulmonary basal, bronchial and nasal epithelial cells

We harvested ‘healthy’ pulmonary epithelial basal cells to model class I nonsense *CFTR* p.(GlyG542Ter), usually named G542X, responsible for cystic fibrosis (NM_000492.4(*CFTR*): c.1624G>T, Fig. 3a). Cells were edited with UNZIP or ZIPmax by nucleofection. The template contained i) a non-coding TAG codon to replace the GGA codon (Gly) and ii) a *SacI* restriction site to allow HDR editing analysis by RFLP and mute the PAM to avoid a Cas9 recut. Analysis revealed a 3-fold higher percentage of HDR-edited alleles with the G542X mutation and the other four nucleotides responsible for the *SacI* cut with ZIPmax compared to UNZIP, reaching up to 22% (Fig. 3b).

**Fig. 3:**
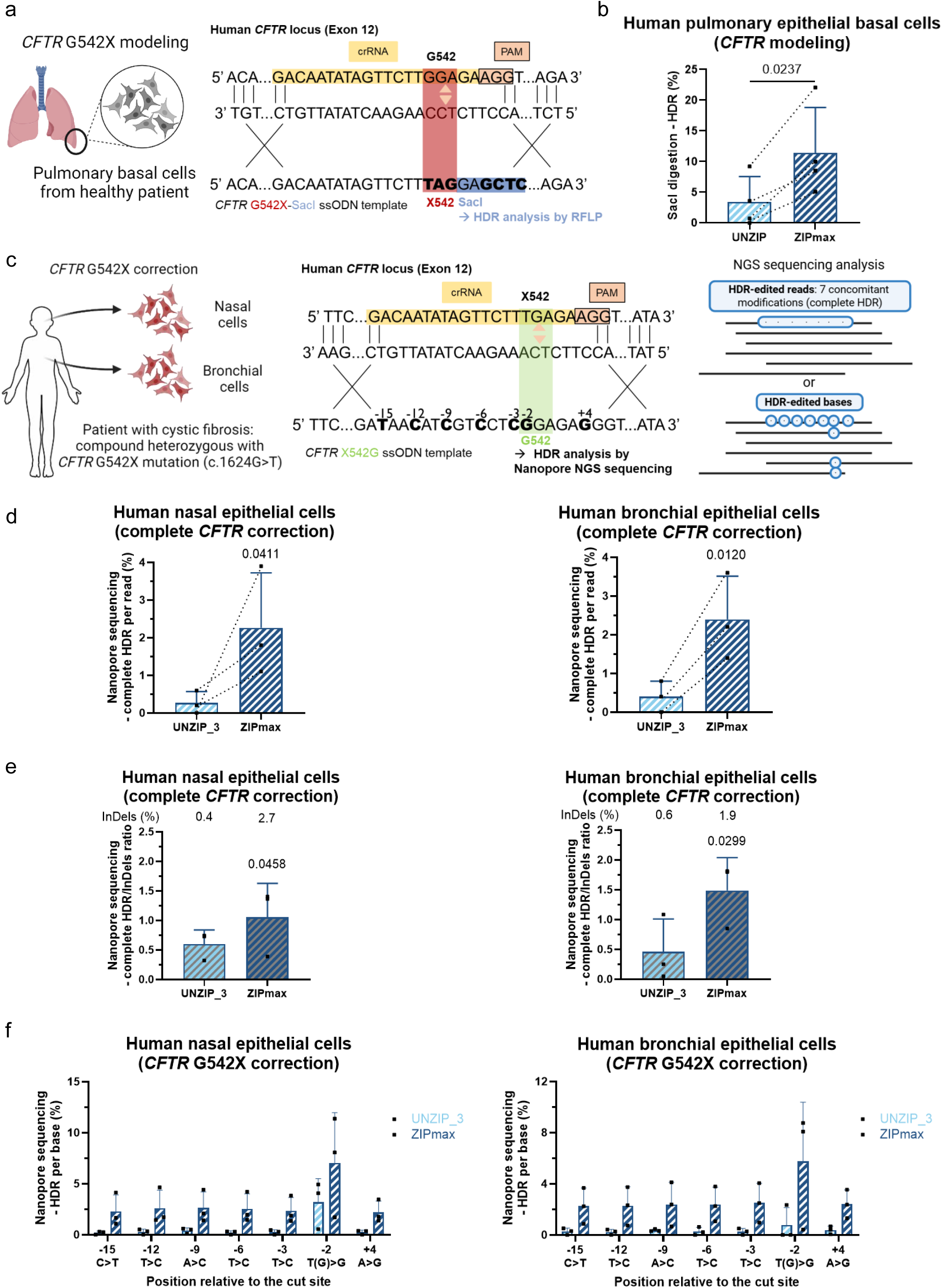
Validation of ZIP CRISPR technology for *CFTR* editing in pulmonary basal, bronchial and nasal epithelial cells. Obtention of human healthy pulmonary epithelial basal cells and editing strategy to model *CFTR* G542X mutation and insert *SacI* restriction site. **b** Capillary electrophoresis of *SacI* digestion products to quantify HDR editing with UNZIP or ZIPmax (n=4). The mean ± SD is shown. Statistical significance determined by paired t-test. **c** Obtention of human nasal and bronchial epithelial cells from a patient with cystic fibrosis carrying the *CFTR* G542X rare mutation and editing strategy i) to correct *CFTR* G542X mutation and ii) to add other silent modifications to prevent re-cut by nuclease Cas9. **d** Nanopore sequencing to quantify HDR editing per read (complete HDR, 7 modifications) with UNZIP_3 or ZIPmax in nasal (left panel) or bronchial (right panel) epithelial cells (n=3). The mean ± SD is shown. Statistical significance compared to unedited cells determined by one-way ANOVA. **e** Nanopore sequencing analysed with nCRISPResso2 to quantify HDR/InDels ratio with UNZIP_3 or ZIPmax in nasal (left panel) or bronchial (right panel) epithelial cells (n=3). The mean ± SD is shown. Statistical significance compared to unedited cells determined by Kruskal Wallis test. The InDels rate is indicated at the top of the graph. **f** Nanopore sequencing to quantify HDR editing per base with UNZIP_3 or ZIPmax in nasal (left panel) or bronchial (right panel) epithelial cells (n=3). The mean ± SD is shown. Schemas created with BioRender.com.

We then decided to correct this nonsense mutation, which is not eligible for current pharmacological treatments^30^. We obtained nasal and bronchial epithelial cells from a cystic fibrosis patient with the compound heterozygous G542X mutation. For editing, we used a mutation-specific gRNA targeting the G542X-mutated allele to keep the other allele (not mutated at this exon) intact, and a correcting template coding GGA (Gly) instead of TGA (nonsense). We added six additional silent mutations (from –15 to +4nt around the cut site) to avoid the Cas9 recut and analyzed editing by Nanopore NGS sequencing (Fig. 3c). Whereas the UNZIP_3 system was inefficient to edit these nasal and bronchial epithelial cells, ZIPmax allowed moderate complete (7 modifications) HDR editing (Fig. 3d left and right panels, around 2%) and increased the HDR/InDels ratio (Fig. 3e). Regarding the c.1624T>G correction specifically, cells were initially 50% c.1624G. We obtained around 56 or 57 % of c.1624G after editing, i.e. 6 or 7% of genomic edition in bronchial and nasal epithelial cells, respectively (Fig. 3f). This result, which is higher than for other substitutions, suggests an additive pathway to correct c.1624T, in accordance with previous reports of gene conversion with the endogenous allele in the event of heterozygosity^31,32^. These HDR editing rates could appear low but have to be put into perspective with the overall editing efficiency and to levels needed to get a clinical improvement. Moreover, regarding the literature, these HDR editing rates (6-7% considering only the c.1624T>G correction relevant for therapeutic impact) could be enough to recover CFTR function and lead to clinical benefit for the patients^33,34^.

### ON-target genotoxicity outcomes of ZIP CRISPR

Cas9 nuclease is known to induce DSB-dependent ON-target genotoxicity^35,36^. Some recently developed HDR boosters, such as AZD7648, have been shown to exacerbate ON-target genotoxicity, causing frequent kilobase– and megabase-scale deletions, chromosome arm loss, and translocations^28,29^. We therefore investigated whether ZIPmax could similarly modulate this toxicity. To test this hypothesis, we used our previously published fluorescence-assisted megabase-scale rearrangements detection (FAMReD) system that sensitively detects and quantifies megabase-scale loss-of-heterozygosity^37^ (Supplementary Fig. S5a). While increasing HDR editing (Supplementary Fig. S5b), ZIPmax did not increase DSB-dependent megabase-scale LOH frequency compared to UNZIP_3 or to gRNA without template (Supplementary Fig. S5c). Thus, the ZIPmax strategy appeared to be safer than protocols modulating DNA repair pathways. Finally, ZIP CRISPR using Cas9 nuclease increased the PGE rate in many *loci* and cell types, but InDels and genotoxicity were still present. We thus applied ZIP CRISPR to a DSB-free Cas9 (single Cas9^D10A^ nickase), which is known to allow HDR editing but only using plasmid, not RNP transfection^8,16,38–47^.

### ZIP CRISPR coupled with Cas9 nickase to obtain precise genome editing

To edit cells without DSB, we applied ZIPmax with the t-lock element in the gRNA and a 5’-extended (120nt) ssODN template to Cas9 nickase. The Cas9^D10A^ nickase was preferred as the unmodified HNH domain has higher activity^48^ and creates less DSBs^49^. This new system was named ZIPmax_Nick (Fig. 4a). As expected, UNZIP with nickase (named UNZIP_Nick) with free BFP-ssODN template did not edit eGFP^+^-HEK293T cells, eGFP^+^-K562 cells and eGFP^+^-hFFs or only at very low levels (Fig. 4b), confirming previous reports showing that it is challenging to edit the genome using a unique nick^39,50^. Importantly, ZIPmax_Nick induced a 3-fold increase in PGE (BFP^+^ cells) in the eGFP^+^-HEK293T cell line (Fig. 4b, left) and unlocked PGE in the eGFP^+^-K562 cell line (17-fold increase, Fig. 4b middle) and in primary eGFP^+^-hFFs (35-fold increase, Fig. 4b right).

**Fig 4:**
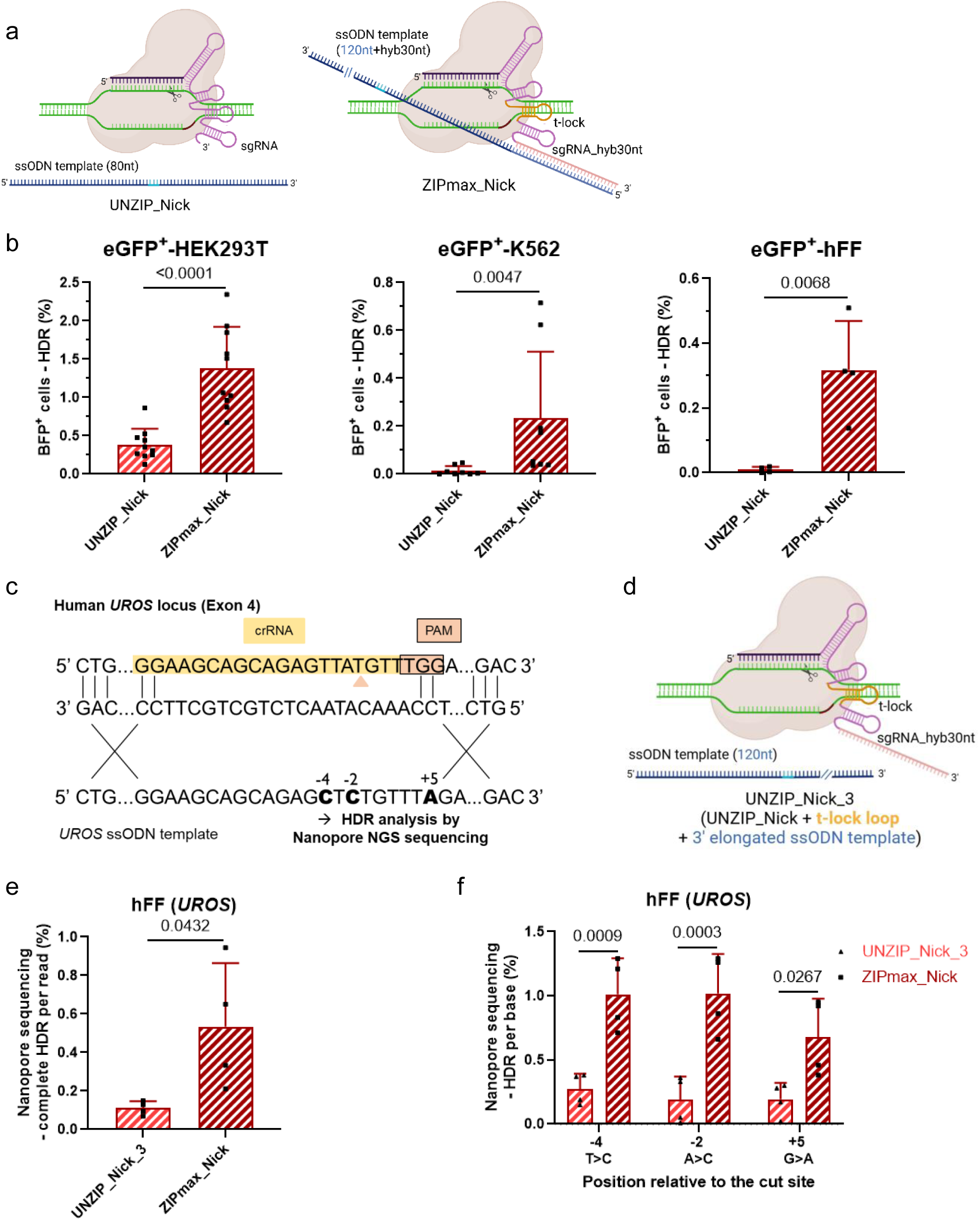
Increased availability of HDR template with ZIPmax_Nick is essential to repair nicks by HDR. **a** Representation of RNPs UNZIP_Nick (Cas9 nickase with free ssODN template) and ZIPmax_Nick (Cas9 nickase with ssODN import using hybridization of 5’-elongated ssODN to the t-lock gRNA). **b** Flow cytometry quantification of the edition with Cas9 nickase with (ZIPmax_Nick) or without (UNZIP_Nick) the correction template physically bound to the gRNA in eGFP^+^-HEK293T cells (n=10), eGFP^+^-K562 cells (n=8), and eGFP^+^-hFFs (n=4). The proportions of cells expressing BFP (HDR-edited) one week after transfection are reported. The mean ± SD is shown. Statistical significance determined by unpaired t-test (eGFP^+^-HEK293T cells, eGFP^+^-hFFs) or Mann-Whitney test (eGFP^+^-K562 cells). **c** Editing strategy to edit *UROS* in hFFs. **d** Representation of UNZIP_Nick_3 (control of ZIPmax_Nick with modified gRNA but without ssODN-gRNA annealing). **e** Nanopore sequencing to quantify HDR editing per read (complete HDR, 3 modifications) with UNZIP_Nick_3 or ZIPmax_Nick (n=4). The mean ± SD is shown. Statistical significance determined by unpaired t-test. **f** Nanopore sequencing to quantify HDR editing per base with UNZIP_Nick_3 or ZIPmax_Nick (n=4). The mean ± SD is shown. Statistical significance determined by two-way ANOVA. Schemas created with BioRender.com.

To confirm these results, we edited an endogenous *locus* (*UROS*) in hFFs with a template containing three substitutions (Fig. 4c). To be sure that the difference between UNZIP_Nick control and ZIPmax_Nick was really due to the import system and not to other modifications (t-lock or the 3’-extension in the gRNA), we designed a new control with Cas9^D10A^ nickase similar to UNZIP_3 using nuclease with a gRNA containing the t-lock element and the 3’-extension, but with a free 120nt template (without 5’-extension, UNZIP_Nick_3, Fig. 4d). Nanopore sequencing revealed a 4-fold improvement of perfect *UROS* HDR editing (3 concomitant edited bases, Fig. 4e), confirmed by individual base analysis (Fig. 4f) and without any detection of InDels (not shown).

Unlike for the nuclease, the absence of InDels using the nickase was an opportunity to repeat editing of the unedited alleles using the same gRNA to enrich the HDR-edited population (Fig. 5a). We made three iterative RNP transfections with ZIPmax_Nick in eGFP^+^-hFFs and obtained an interesting additive effect leading to more than 3% of BFP^+^ cells (Fig. 5b). We confirmed this additive effect targeting *UROS* in hFFs (Fig. 5c).

**Fig. 5:**
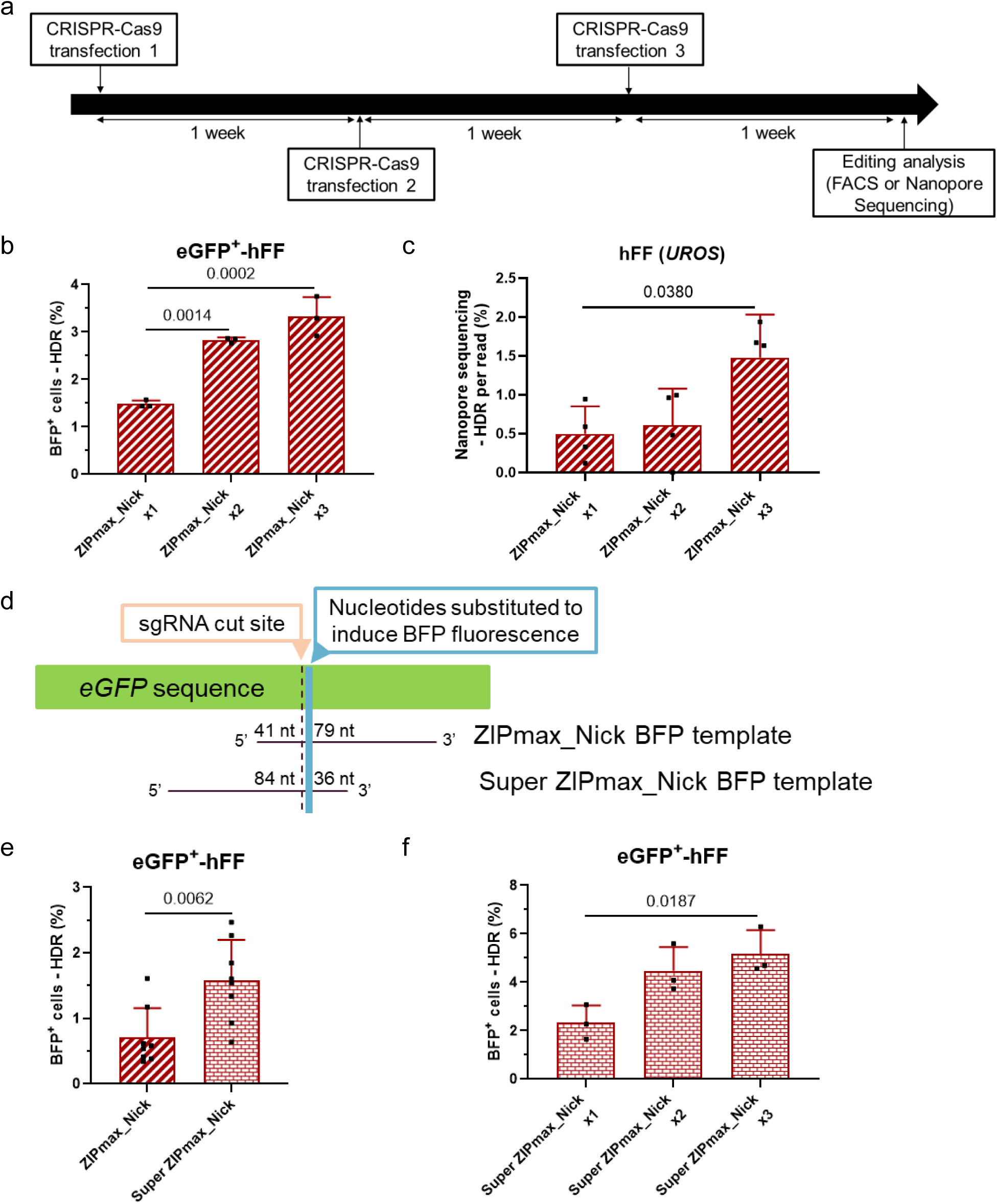
Iterative editing and ssODN template design improve HDR editing with ZIPmax_Nick. **a** Timeline for iterative transfections (RNPs nucleofection). **b** Flow cytometry quantification of edition after 1, 2 or 3 transfection(s) with ZIPmax_Nick in eGFP^+^-hFFs (n=3). The proportions of cells expressing BFP (HDR-edited) one week after transfection are reported. The mean ± SD is shown. Statistical significance determined by one-way ANOVA. **c** Nanopore sequencing quantification of edition after 1, 2 or 3 transfection(s) with ZIPmax_Nick in hFFs (n=4). Complete HDR (3 modifications) rates per read are reported. The mean ± SD is shown. Statistical significance determined by one-way ANOVA. **d** Representation of BFP-ssODN template position relative to the cut site used for ZIPmax_Nick or Super ZIPmax_Nick. **e** Flow cytometry quantification of edition with ZIPmax_Nick or Super ZIPmax_Nick in eGFP^+^-hFFs (n=8). The proportions of cells expressing BFP (HDR-edited) one week after transfection are reported. The mean ± SD is shown. Statistical significance determined by unpaired t-test. **f** Flow cytometry quantification of edition after 1, 2 or 3 transfection(s) with Super ZIPmax_Nick in eGFP^+^-hFFs (n=3). The proportions of cells expressing BFP (HDR-edited) one week after transfection are reported. The mean ± SD is shown. Statistical significance determined by one-way ANOVA.

Then, to take advantage of our easy-to-design ZIP CRISPR tool, we evaluated the impact of the ssODN design and compared two asymmetric templates with either the nucleotide modifications on the 5’-(ZIPmax_Nick template) or on the 3’-extremity (Super ZIPmax_Nick template) (Fig. 5d). Interestingly, we observed a 2-fold increase in HDR editing in eGFP^+^-hFFs with Super ZIPmax_Nick compared to ZIPmax_Nick (0.71% ± 0.45 *vs* 1.58% ± 0.62 respectively, Fig. 5e). Thus, ZIP CRISPR could be a great help to test and optimize template designs. With Super ZIPmax_Nick in eGFP^+^-hFFs, we confirmed that iterative transfections led to an additive effect and resulted in more than 5% of HDR-edited cells (Fig. 5f).

### ZIPmax_Nick can be combined with ‘cell state’ modulation to increase nick-based HDR editing

To further improve HDR editing efficiency and benefit from the versatility of ZIPmax_Nick, we investigated various pathways known to modulate genome editing. First, we tried to modulate DNA repair pathways. Nick-mediated HDR repair using a single-stranded template is known to be suppressed by RAD51/BRCA2^39^. Thus, we transiently inhibited *BRCA2* using a siRNA in eGFP^+^-HEK293T cells and eGFP^+^-K562 cells. In both cell lines, we obtained a transient drop in *BRCA2* mRNA expression (Supplementary Fig. S6a). We achieved a 2-fold increase in HDR editing (BFP^+^ cells) when BRCA2 was inhibited during editing (Supplementary Fig. S6b). However, it was accompanied by a 3 to 4-fold increase in imprecise editing (InDels, eGFP^-^/BFP^-^ cells) in eGFP^+^-HEK293T cells and eGFP^+^-K562 cells (Supplementary Fig. S6c), probably due to the conversion of nicks into DSBs. Thus, we decided not to pursue the idea of inhibiting *BRCA2* to avoid DSB-associated genotoxicity. We also tried to inhibit *MLH1* by a CRISPR knock-out to modulate the mismatch repair (MMR) pathway, but we encountered toxicity issues after editing (data not shown).

While HDR editing using the Cas9 nuclease is known to depend on the S/G2 phases, the impact of the cell cycle on nick-mediated HDR editing has been less studied^51^. We observed higher HDR editing rates in highly proliferative HEK293T cells than in less proliferative hFFs (Fig. 4b), suggesting that editing could preferentially be performed during the S/G2/M phases. To decipher the role of the cell cycle on HDR-based nick repair, and the S/G2/M phases in particular, we modulated hFFs’ cell cycle. We used palbociclib, a CDK4/6 inhibitor, to reduce the proportion of eGFP^+^-hFFs in the S/G2/M phases during editing (Fig. 6a/b). We observed a 2-fold decrease in HDR editing rates (Fig. 6c and Supplementary Fig. S7a). In contrast, HDR editing efficiency was improved by almost 2-fold (+85.4%, Fig. 6c) when using the optimal conditions of CDK7 inhibitor XL413 during editing (Supplementary Fig. S7b/c/d/e/f) to increase the proportion of eGFP^+^-hFFs in S/G2/M phases (Fig. 6a/b). These data suggest that HDR editing with nickase is S/G2/M phase-dependent.

**Fig. 6:**
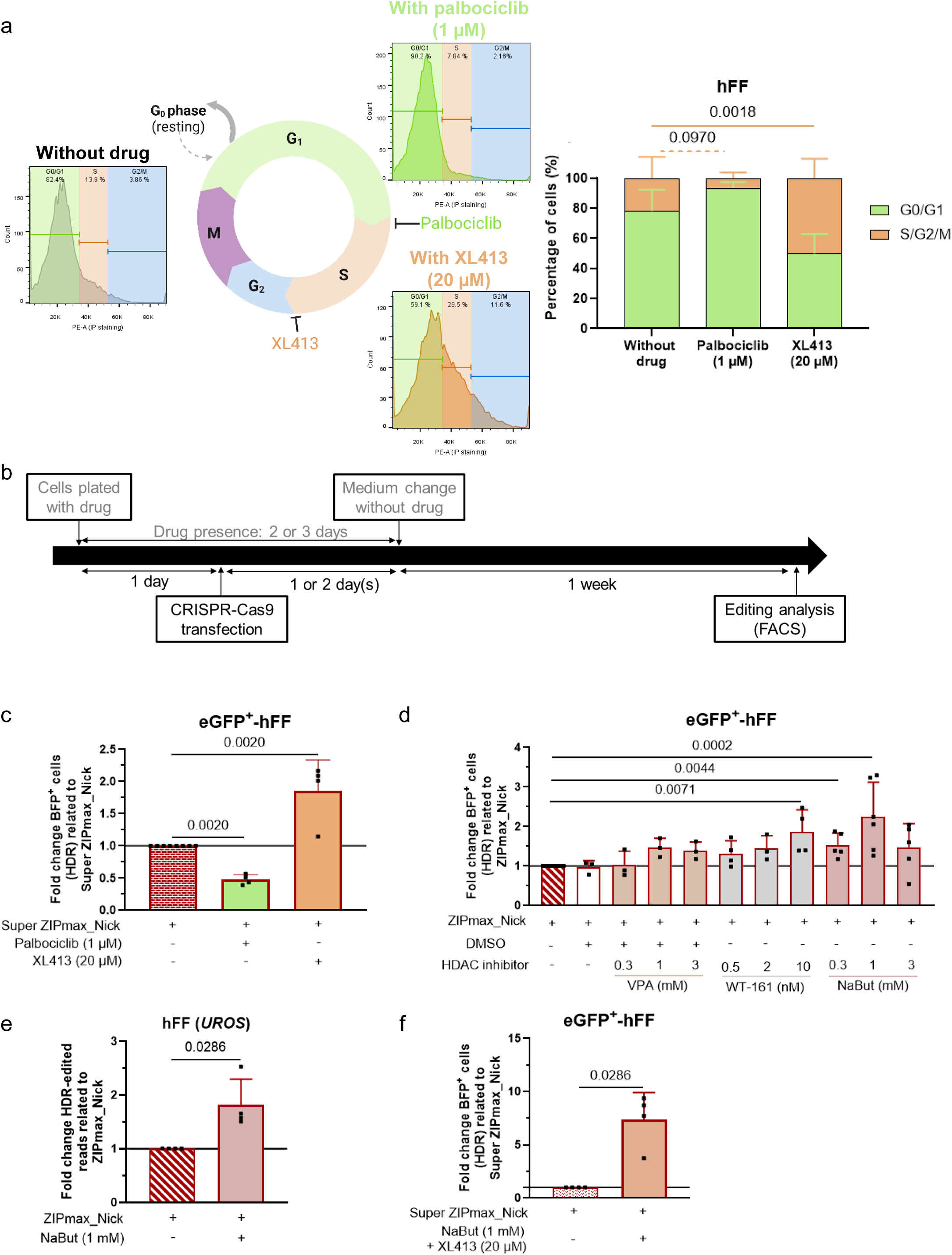
Modulation of ‘cell state’ can be associated with ZIPmax_Nick to increase HDR editing. **a** Illustrative cell cycle results of eGFP^+^-hFFs with palbociclib, XL413 (right panel) or without drug (left panel). Flow cytometry quantification of the proportion of eGFP^+^-hFFs in G0/G1 or S/G2/M phases with palbociclib, XL413 or without drug (right panel, n=4). The mean ± SD is shown. Statistical significance to cells without drugs determined by two-way ANOVA. **b** Timeline for transfections (RNPs nucleofection) in presence of drugs. **c** Flow cytometry quantification of HDR edition with Cas9 nickase (Super ZIPmax_Nick) with or without synchronization of eGFP^+^-hFFs in G0/G1 phase with palbociclib or in S/G2/M phases with XL413 (n=4). The proportions of cells expressing BFP (HDR-edited) one week after transfection are reported. Fold changes indicate fold increases between conditions with or without drug. The mean ± SD is shown. Statistical significance to Super ZIPmax_Nick without drugs determined by Kolmogorov-Smirnov test. **d** Flow cytometry quantification of HDR edition with nickase Cas9 (ZIPmax_Nick) with or without VPA (n=3), WT-161 (n=3) or NaBut (n=5) exposure at different doses. The proportion of cells expressing BFP (HDR-edited) one week after transfection is reported. Fold changes indicate fold increases between the conditions with either VPA, WT-161 or NaBut and the condition without drug. The mean ± SD is shown. Statistical significance determined by Kruskal-Wallis test. **d** Nanopore sequencing to quantify *UROS* HDR editing per read with Cas9 nickase (ZIPmax_Nick) with or without NaBut in hFFs (n=4). Fold changes indicate fold increases between the conditions with or without NaBut. The mean ± SD is shown. Statistical significance determined by Kolmogorov-Smirnov test. **f** Flow cytometry quantification of HDR edition with Cas9 nickase (Super ZIPmax_Nick) with or without NaBut and XL413 exposure in eGFP^+^-hFFs (n=4). The proportions of cells expressing BFP (HDR-edited) one week after transfection are reported. Fold changes indicate fold increases between conditions with or without drugs. The mean ± SD is shown. Statistical significance to Super ZIPmax_Nick without drugs determined by Kolmogorov-Smirnov test. Schemas created with BioRender.com.

Chromatin access can also be important for the efficiency of CRISPR tools. We modulated chromatin opening by using HDAC inhibitors in eGFP^+^-hFFs with ZIPmax_Nick. When we screened several drugs (valproic acid – VPA –, WT-161 and sodium butyrate – NaBut –) at different doses, we observed a global trend to an increase in HDR editing efficiency in the presence of HDAC inhibitors. A significant effect was found with WT-161 at 10 nM (+86.6%), NaBut at 0.3 mM (+52.7%) and more importantly with NaBut at 1 mM (+125%) (Fig. 6d and Supplementary Fig. S7g/h). We validated the effect of NaBut at 1 mM editing *UROS* in hFFs (+81.5%, Fig. 6e and Supplementary Fig. S7i). Then, we combined NaBut and XL413 exposure. Remarkably, we observed an additive effect, leading to a 7-fold increase in HDR editing compared to the condition without drugs (Fig. 6f and Supplementary Fig. S7j).

Therefore, by reaching detectable HDR editing rates at several *loci* and in several cell types with ZIPmax_Nick, we demonstrated that HDR editing with nickase was more efficient during or after replication, and was boosted by HDAC inhibitors and ssODN template design.

## Discussion

HDR editing remains a challenge with CRISPR-Cas9, especially for clinical purposes. The spatiotemporal availability of the HDR template at the cut site during repair is supposed to be a key factor for HDR editing. In this study, we explored the import capacity of annealing the ssODN HDR template to the gRNA to increase PGE through DSBs or nicks. Using RNPs, we demonstrated that ZIP CRISPR is an ssODN import system easy to implement that considerably increases HDR editing efficiency. FRET demonstrated the increased proximity of ssODNs and RNP complexes and the stability of the complex into the transfected cells. Moreover, FRET experiments suggest that part of this increase could also stem from enhanced transfection or stability of the gRNA–ssODN complex upon annealing. We further optimized ZIP CRISPR for several targets (*eGFP*, *HBB*, *UROS*, *CFTR*, *SORCS1*) in different cell types (HEK293T cells, K562 cells, hFFs, HSPCs and pulmonary, bronchial and nasal epithelial cells) and for various kinds of edits (substitutions of 2 to 7 bases and deletions) around the cut site (editing window from –15 to +15nt depending on the targets). We obtained up to 12-fold improvement in HDR editing when mediated by a DSB (mean of 5-fold) and up to 35-fold (unlocking) when mediated by a single nick compared to a control approach with a free ssODN template.

As previously described in the introduction, other strategies have been implemented to bring the ssODN with the Cas9/gRNA complex, with a link to the Cas9 or the gRNA, but faced some limitations especially in terms of production, transfection, gene therapy immunogenicity, and the increased costs of producing a gene-editing tool. Here we develop a chemical modification-free approach that can recruit ssDNA donors to the target site and enhance HDR efficiency. Unlike the other approaches, ZIP CRISPR is free from fusion protein or a covalent link between CRISPR-Cas9 components. Other strategies were developed to increase the concentration of the ssODN template near the DNA break. Polymerase-based DNA writing technologies have been developed to produce large template quantities *in situ* near the targeted *locus*. Prime editing enables precise modification of genomes *via* reverse transcription of an RNA pre-template encoded at the 3’ end of the gRNA (a prime editor guide RNA, pegRNA) to produce the ssODN template^24^. More recently, a DNA-dependent polymerase-based system known as click editing was developed to amplify the ssODN template with higher fidelity^52,53^. However, these strategies rely on large Cas9 fusion proteins featuring a polymerase activity that still require limiting substrates (nucleotides, MgCl2…). In addition, their cell impact remains largely elusive, and they are complex to design and adapt to new targets.

In this study, we used ZIP CRISPR with ssODN templates shorter than 200nt. However, it would be interesting to use it to import larger single-stranded DNA or double-stranded DNA with a 5’-ssDNA extremity to anneal to the gRNA. This could be a safer non-viral alternative to the AAV or IDLV template currently used for large gene insertions^54^. Further studies would be important to explore this possibility.

Importantly, ZIP CRISPR reached high-efficiency HDR editing rates without modulating DNA repair pathways. Indeed, several methods have been proposed to inhibit NHEJ/MMEJ. Small pharmacological molecules (e.g. SCR7, AZD7648, M3814) have been developed to transiently inhibit the NHEJ components (KU70/80, DNA PK, Ligase IV)^2,3^. The concomitant inhibition of NHEJ and MMEJ by polymerase theta (Polθ) inhibitors has also been proposed^4–6^. In all studies, these molecules limit the NHEJ/MMEJ pathways and therefore enhance HDR editing efficiency. Recruitment^7^ or ectopic expression of RAD51 alone or combined to SCR7^8,9^ can also be used. In the present study, ZIP CRISPR was at least as efficient as AZD7648 to increase the HDR editing rate in hFFs. Interestingly, the efficiency was improved when ZIP CRISPR and AZD7648 were associated. This combined approach could be interesting to obtain higher levels of HDR editing and confirm that the selection of the DSB repair pathway and the recruitment of the DNA template are the two main steps limiting the rate of HDR editing. However, modulating DNA repair pathways may not be without effect and could alter genome integrity. For example, one of the most potent NHEJ inhibitors, AZD7648, was recently shown to be genotoxic^28,29^ inducing large deletions and translocations. This suggests that DNA repair modulators should be used carefully in clinical studies and that researchers risk overestimating the results of HDR rate^36^. Interestingly, ZIP CRISPR did not increase DSB-associated genotoxicity and HDR editing efficiency in hFFs was higher with ZIPmax than with AZD7648 combined with UNZIP. The safety of ZIP CRISPR has to be considered for future clinical applications of CRISPR-Cas9 mediated gene therapy, and ZIP CRISPR should be preferred to NHEJ inhibitors.

To avoid genotoxicity, editing without DSBs is recommended for gene therapy. SSB-based editing with single nickase is feasible but currently relies on plasmid transfection of components to reach detectable HDR-edition levels. This lack of efficiency^38,55,56^ is probably due to the involvement of the single-strand break repair (SSBR)^39^ and MMR repair pathways. Indeed, it has been described for other SSB-based CRISPR tools, such as base editing^57^ and prime editing^58,59^ that editing was higher in MLH1-deficient cells and the authors have proposed to inhibit MLH1 (by knock out or using dominant negative) or to trick the MMR pathway with the introduction of a second nick on the opposite strand. However, these approaches are toxic or induce more InDels and ON-target genotoxicity^60^. Here, ZIP CRISPR allowed us to obtain HDR editing without InDels, with a RNP single nickase (to avoid any risk of genomic integration), and without polymerase or fusion protein. Using this new tool we decipher the mechanisms implicated in nick-mediated HDR-editing.

HDR repair at nicks is known to be influenced by the transcriptional status of the nicked strand (more efficient on the transcribed strand) and suppressed by RAD51/BRCA2 when using an ssDNA template^39^. Here, we confirmed the inhibiting role of BRCA2 on nick-based HDR editing and showed an important increase in imprecise editing, validating BRCA2’s importance in maintaining genome integrity^61^.

We also investigated the impact of cell cycle modulation on nick-based HDR editing, showing that HDR editing was increased in proliferative cells (synchronized in the S + G2/M phases) and reduced in non-dividing cells (synchronized in the G0/G1 phases). Those results are different from a previous study done by Zhang, Davis and Maizels in 2021^51^ showing that nicks initiated HDR more efficiently in the G1 than in the S/G2 phases. This could be explained by the different cells (primary fibroblasts in this study *vs* HEK293T and U2OS cell lines) and the different methodologies used: pharmacological synchronization of cells in a phase of the cell cycle for 48h *vs* the use of Cas9 fused to degrons, as the previous publication indicated that the G1 phase nicks may persist to be repaired later in cell cycle. We also confirm chromatin opening impact for nick-based HDR editing in line with previous studies done with other CRISPR-based editing tools^62,63^. Further studies will be required to improve the efficiency and decipher the nick-based HDR editing mechanisms.

Altogether, ZIP CRISPR is a new programmable chemical-free non-viral versatile tool that can meet the needs of a wide range of experiments. ZIPmax based on nuclease is the first choice when a higher HDR editing rate is needed and when DSB-associated genotoxicity is acceptable. On the other hand, ZIPmax_Nick tools based on single nickase are more suited when a non-genotoxic approach is required and when low HDR editing is sufficient to restore the phenotype, or when iterative editing is feasible. The simplicity of ZIP CRISPR and all its versions should extend their utility to the investigation of complex biological issues in a wide range of cells and organisms. Since ZIP CRISPR is modular, it is potentially compatible with several types of templates and Cas, so it could play a leading role in many areas of research. Importantly, ZIP CRISPR achieves efficient HDR editing and could be considered to unlock non-viral HDR-based gene therapies.

## Methods

### Ethical statement

This research complies with all relevant ethical regulations. Human CD34^+^ stem and progenitor cells (HSPCs) were isolated from cord blood of healthy donors from Bagatelle Hospital, according to the hospital’s ethical institutional review board (Maison de Santé Protestante de Bordeaux, Talence, France) and with the mothers’ informed consent.

The patient with cystic fibrosis was recruited at the pediatric cystic fibrosis resource and skills center (CPP 23.00854.000207). Nasal and bronchial epithelial cells were obtained by nasal or bronchial superficial ciliary brushing, respectively, according to the hospital’s ethical institutional review board and with the parents’ informed consent.

Control subjects (CPP 23.00854.000207) were recruited after surgical resection at the University Hospital of Bordeaux. Bronchial specimens from all subjects were obtained either by fibroscopic bronchoscopy or lobectomy in macroscopically normal areas, as previously described^64^. All subjects gave their written informed consent to participate in the study after the nature of the procedure had been fully explained.

### Cell culture

Human embryonic kidney (HEK) cell line immortalized with large T antigen HEK293T (ATCC®, Manassas, VA, USA) was maintained in Dulbecco’s modified Eagle’s medium (DMEM), low glucose (1 g/L), L-Glutamine (1 g/L) and pyruvate (Gibco® by Life Technologies™, Carlsbad, USA) supplemented with 10% fetal bovine serum, 100 U/mL penicillin and 100 μg/mL streptomycin (all from Eurobio™, Courtaboeuf, France).

K562 cell line (ATCC®) was maintained in Roswell Park Memorial Institute (RPMI) Medium 1640, L-Glutamine, 25 mM HEPES (Gibco® by Life Technologies™) supplemented with 10% fetal bovine serum (Eurobio™), GlutaMAX (Gibco® by Life Technologies™), 100 U/mL penicillin and 100 μg/mL streptomycin (Eurobio™).

hTERT immortalized human foreskin fibroblasts (hFFs, ATCC®) were maintained in DMEM, high glucose (4.5 g/L), L-Glutamine (1 g/L) and pyruvate (Gibco® by Life Technologies™) supplemented with 10% fetal bovine serum (Eurobio™), 1% essential amino acids (Gibco® by Life Technologies™), 100 U/mL penicillin, 100μg/mL streptomycin (Eurobio™), 10 µg/mL ciprofloxacin (Biogaran™, Colombes, France) and 0.5 µg/mL amphotericin B (Sigma-Aldrich®, Saint Louis, USA). A previously established heterozygous *UROS*^+/-^ hFF cell line was used for the FAMReD genotoxicity assay (see ^37^ for details).

Human HSPCs were isolated from the cord blood of healthy donors. Briefly, mononuclear cells were isolated by Ficoll gradients. hCD34^+^ cells were purified according to the manufacturer’s instructions (Human CD34-Positive Selection kit II from Stem Cell Technologies, Vancouver, Canada) and purity was analyzed by flow cytometry using phycoerythrin-conjugated anti-CD34 antibody (Biolegend, San Diego, USA). Cryopreserved hCD34^+^ cells were thawed and cultured in expansion medium consisting in StemSpan SFEM (Stem Cell Technologies) supplemented with Flt3-L (100 ng/mL, Peprotech, Carlsabad, USA), SCF (100 ng/mL Peprotech), human TPO (100 ng/mL Peprotech), vitamin C (0.35 mg/mL, Sigma-Aldrich), ciprofloxacin (10 µg/mL, Biogaran™) and 100 U/mL penicillin and 100 μg/mL streptomycin (Eurobio™).

Nasal and bronchial epithelial cell cultures were established from nasal or bronchial brushings, as previously described^64,65^. Nasal and basal epithelial cells were cultured in PneumaCult™-Ex Plus Medium (StemCell Technologies) supplemented with penicillin-streptomycin-Amphotericin B 1X (100X, Gibco).

All cell types were cultured at 37 °C, 5% CO2 in a humidified chamber.

### Construction of eGFP^+^ cell models

HEK293T cells, K562 cells and hFFs were transduced with a lentivirus containing the gene encoding eGFP (enhanced green fluorescent protein) and a puromycin resistance gene, under the dependence of the ubiquitous strong constitutive promoter MND (myeloproliferative sarcoma virus enhancer, negative control region deleted, dl587rev primer-binding site substituted). This lentivirus was produced and titrated by the Vect’UB vectorology platform at Bordeaux University. HEK293T cells, K562 cells and hFFs were transduced at a multiplicity of infection (MOI) of 0.1, 1 and 2, respectively, to obtain a low copy number of the integrated vector. Transduced cells were selected 96h after transduction by adding puromycin (2 μg/mL for 48h, Gibco) to their culture medium. Hereafter, these cells are referred to as eGFP^+^-HEK293T cells, eGFP^+^-K562 cells and eGFP^+^-hFFs.

### RNP transfection and gene-editing tools

The crRNAs were designed using CHOP-CHOP software (chopchop.cbu.uib.no). Then the tracrRNA part (eventually modified for one of the loops and lengthwise) was added to reconstitute the gRNA. gRNAs and ssODN templates for HDR correction were ordered from Integrated DNA Technologies (Coralville, USA). Their sequences are given in Supplementary Tables 1, 2 and 3.

Eventually, annealing between sgRNA and ssODN template(s) to form sgRNA-template hybrids (ZIP, ZIP_Lock, ZIP_Long, ZIPmax, ZIP_Long_Lock_hyb15, ZIP_Long_Lock_hyb20, ZIP_ExtraLong_Lock, ZIP_Inv, ZIPmax_V2, RAD51_ZIPmax, ZIPmax_RAD51, ZIPmax_Nick, Super ZIPmax_Nick) or sgRNA-templates hybrid (ZIP_Double_Lock) was performed by mixing the components either in equimolar proportion (1:1) or with a double template (1:2) for the ZIP_Double_Lock. The components were then heated for 5 minutes in a dry bath at 95°C before allowing them to cool at room temperature.

The different components of the CRISPR-Cas9 system (Alt-R® S.p. HiFi Cas9 Nuclease V3 (abbreviated hereafter as Cas9 nuclease) or Alt-R S.p.Cas9 D10A Nickase V3 protein (abbreviated hereafter as Cas9 nickase) both from Integrated DNA Technologies, with either split sgRNA and ssODN template or sgRNA-template(s) hybrid(s) were combined to form an RNP. They were mixed and then incubated for 10-20 minutes at room temperature. For this, 62 pmol (for HEK293T cells, K562 cells and hFFs) or 105 pmol (for other cells) of nuclease or nickase Cas9 and 100 pmol (for HEK293T cells, K562 cells and hFFs) or 170 pmol (for other cells) of sgRNA, ssODN template or sgRNA-template(s) hybrids for ZIP CRISPR were mixed. Finally, 100 pmol of Alt-R Cas9 Electroporation Enhancer solution (Integrated DNA Technologies) were added to the RNP complex to improve electroporation efficiency.

For S1mplex transfection, S1m-gRNA and biotin-ssODN were ordered from Integrated DNA Technologies. The RNP complex was formed as described in the publication^22^. Molecular ratios were conserved but quantities were adapted to use the same quantity of Cas9 as in the other conditions of the experiment.

Cells were transfected by electroporation using the Nucleofector 4D AMAXA electroporation system (Lonza®, Bale, Switzerland). In brief, 5.10^4^ to 3.10^5^ depending on the cell type were resuspended in SF Cell Line 4D-Nucleofector® (HEK293T cells or K562 cells) or P3 Primary Cell Line 4D-Nucleofector® (hFFs, CD34^+^ HSPCs, pulmonary epithelial basal cells, nasal epithelial cells and bronchial epithelial cells) and added to the RNP complex. Then cells were nucleofected using DG-150, FF-120, CZ-167, DO-100 or DC-100 programs, respectively.

Cells were then cultured as described above for 5-15 days depending on the editing analysis method.

### Edition analysis

#### HDR and imprecise editing (InDels) analysis by flow cytometry

For the *eGFP* target, precise editing allows the switch from eGFP to BFP. At least 6 days after editing and before flow cytometry analysis, cells were recovered in a PBS (phosphate buffered saline, Gibco) – EDTA (ethylenediaminetetraacetic acid, 2 mM, Sigma-Aldrich) solution. Cells were analyzed for fluorescence using the BD FACSCanto II analyzer (BD Biosciences, Le Pont de Claix, France) at the cytometry platform at University of Bordeaux (UB’FACSility). Excitation wavelengths of 488 and 405nm were used to detect eGFP and BFP fluorescence in the FITC (Fluorescein-5-isothiocyanate) (530/30 filter) and Pacific Blue (450/50 filter) channels, respectively. The data presented are from the analysis of at least 10,000 events. The percentages of eGFP^+^/BFP^-^, eGFP^-^/BFP^+^, and eGFP^-^/BFP^-^ correspond to cells unedited, edited by HDR, or imprecisely edited, respectively. The percentages of BFP^+^ or eGFP^-^/BFP^-^ cells shown in the graphs were calculated by subtracting the percentages of BFP^+^ or eGFP^-^/BFP^-^ cells, respectively, from a sample of cells from the corresponding lineage not transfected with RNP when these were non-zero. Then, the percentage of BFP^+^, eGFP^-^/BFP^-^ or eGFP^+^/BFP^-^ cells were normalized to recover a total of 100%.

#### HDR analysis by RFLP

If a restriction site is added in the ssODN template sequence, the HDR rate can be measured by RFLP (restriction fragment length polymorphism). At least 5 days after editing, genomic DNA was extracted from cell pellets using Nucleospin® Tissue (Macherey-Nagel, Duren, Germany) according to the manufacturer’s protocol. The genomic region flanking the edited site was amplified by PCR (HotStarTaq Plus DNA polymerase, Qiagen®, Venlo, Netherlands) with adequate primers (Supplementary Table 4). PCR products were purified with Nucleospin® Gel and PCR Clean-up (Macherey-Nagel) and digested with SacI restriction enzyme (New England Biolabs, Ipswich, USA) for at least 1 h at 37 °C. Then, digestion products were loaded into the Agilent® 2200 TapeStation (Santa Clara, USA) capillary electrophoresis using D1000 ScreenTape and D1000 reagents according to the manufacturer’s protocol. Quality control of enzymatic digestion efficiency was included in each assay.

#### HDR and InDels analysis by Nanopore sequencing

At least 5 days after editing, genomic DNA was extracted from cell pellets using Nucleospin® Tissue (Macherey-Nagel) according to the manufacturer’s protocol. The genomic region flanking the edited site was amplified by High-Fidelity PCR (Phusion High-Fidelity DNA Polymerase, New England Biolabs) using 35 cycles under recommended manufacturer’s protocol with adequate primers (Supplementary Table 4). A sequencing library was prepared using the Rapid Barcoding Kit 96 V14 (SQK-RBK114.96, Oxford Nanopore Technologies, Oxford, UK) according to the manufacturer’s protocol. Sequencing was then performed using a MinION Mk1B device and a Flow Cell R10.4.1 (FLO-MIN114) both from Oxford Nanopore Technologies to reach at least 2000 quality-filtered reads per sample (Q score > 8). After sequencing, super-accurate basecalling was performed using MinKNOW (v24.02.6) and FASTQ were aligned on the human reference genome GrCH38 (GCA_0000001405.15) using Epi2Me (v5.1.9) and the wf-alignment pipeline (v1.1.2). Then, BAMs were analyzed with two custom bioinformatic scripts in Python (v3.8) using pysam (v0.16). Briefly, one searches specific sequences in the BAM corresponding to unedited, HDR edited sequences or eventually predominant InDels seen with IGV (Integrative Genomics Viewer) visualization to determine the HDR editing rate per read, whereas the other one calculates the proportion of each nucleotide for each selected position to determine the HDR editing rate per base, returning these data in xlsx format (pandas v1.1.3). For *CFTR* G542X correction, HDR and InDels rates were determined using nCRISPResso2, according to Mc Farlane et al’s publication^66^. The command is available on Supplementary Table 5.

### Drugs used

#### Cell cycle synchronization

To evaluate the impact of synchronization on HDR editing efficiency, cells were synchronized in the G0/G1 phase by incubation with palbociclib, also named PF-00080665 or PD 0332991 (1 µM, Sigma Aldrich) 24 h before and 24 h after RNP transfection. To synchronize cells in the S phase, they were incubated with XL413 (10, 20 or 40 µM, Selleckchem, Houston, USA) for 24 h before RNP transfection and 24h after RNP transfection (or at the time indicated in the figures).

Cell cycle analyses were performed to check synchronization efficiency in each experiment. Briefly, cells either synchronized or not were harvested, fixed with 4% paraformaldehyde (for 15 minutes) and permeabilized with Triton 0.5% (Sigma-Aldrich, for 15 minutes). Cells were then incubated overnight with RNAse (100 µg/mL, MoBiTec, Goettingen, Germany) and then for 15 minutes with propidium iodide (4 μg/mL, Biolegend). The samples were examined on a BD Biociences Accuri C6 Plus flow cytometer and the data were analyzed with BD CSampler™ software (BD Biosciences).

#### HDAC inhibition

To evaluate the impact of HDAC inhibition, cells were incubated 24h before and 48h after edition (or at the time indicated in the figures) with either valproic acid (VPA, 0.3, 1 or 3 mM, Sigma-Aldrich), WT-161 (0.5, 2 or 10 nM, MedChemExpress, Monmouth Junction, USA) or sodium butyrate (NaBut, 0.3, 1 or 3 mM, Sigma-Aldrich).

#### NHEJ inhibition

To modulate the NHEJ pathway, we incubated the cells 3h before and 48h after edition (or at the time indicated in the figures) with AZD7648 (0.3, 1, 3 or 10 µM, MedChemExpress).

#### Nick repair pathway inhibition

To modulate the nick repair pathways, HEK293T and K562 cells were transfected either 48h or 72h before edition with a siRNA targeting *BRCA2* (10 pmol/mL, #s2085 Life Technologies, Carlsbad, USA), respectively. siRNA transfections were performed using Lipofectamine RNAiMAX (Life Technologies) according to the manufacturer’s protocol. To determine the efficiency of siRNA knockdown, *BRCA2* mRNA levels were quantified by RT-qPCR relative to control that assayed glyceraldehyde-3-phosphate dehydrogenase (*GAPDH*) transcripts. Briefly, RNA was extracted from cell pellets using Nucleospin® RNA (Macherey-Nagel) according to the manufacturer’s protocol. RNA was retro-transcribed into cDNA using high-capacity cDNA reverse transcription (Applied Biosystems, Waltham, USA) according to the manufacturer’s protocol. Then qPCR was performed using GoTaq qPCR Master Mix 1X (Promega, Madison, USA) and adequate primers (Supplementary Table 6). Results were analyzed with CFX Maestro 1.0 (BioRad, Hercules, USA).

### FAMReD fluorescent cell quantification

At least 15 days after editing, 0.3 mM of 5-aminolevulinic acid hydrochloride (5-ALA, Sigma-Aldrich) was added to hFFs’ medium. After 16h of exposure (overnight), cells were washed twice with PBS and placed in fresh medium. Upon LOH, loss of UROS function can be detected by the appearance of fluorescence due to porphyrin accumulation. Following 8h of clearance, fluorescent cells were quantified by flow cytometry. UV-sensitive porphyrins were excited at 488 nm and the emitted wavelength was approximately 667 nm, detected by the PE-Cy5A PMT channel (FACS Accuri, BD Biosciences). FL-1 is a control green-fluorescent channel used to exclude auto-fluorescent cells.

### FRET experiments

Cy5-probe complementary to the sgRNA and Cy3-ssODN template, were designed according to Li Y. *et al.* publication^67^ and ordered from Integrated DNA Technologies. Their sequences are given in Supplementary Table 7. Annealing between sgRNA (with or without hybridization sequence for the ssODN template), Cy5-probe and Cy3-ssODN template was performed by mixing the components in equimolar proportion (1:1). The components were then heated for 5 minutes in a dry bath at 95°C before allowing them to cool at room temperature. Then, Cas9 was added to form an RNP.

For spectrofluorometry analysis, 93 pmol of Cas9 was added to the annealing containing 150 pmol of sgRNA, 150 pmol of Cy5-probe and 150 pmol of Cy3-ssODN template. Components were mixed and incubated for 10 min before analysis. FRET analysis was performed with Hitachi F-4500 fluorescence spectrophotometer using an excitation length of 554 nm (Cy3 excitation length). Relative quantification was done measuring excitation peak height difference at 665 nm (Cy5 emission length) between UNZIP and ZIP conditions.

For confocal microscopy analysis, 62 pmol of Cas9, and depending on the conditions 100 pmol of sgRNA, 100 pmol of Cy5-probe and/or 100 pmol of Cy3-ssODN template were mixed and incubated for 10-20 min before transfection as previously described. For the “low dose” experiment, quantities were divided by three. After transfection, cells were put on slides for 7h (adhesion time). Then medium was withdrawn and cells were fixed with 4% paraformaldehyde (for 15 minutes) and washed three times with PBS. Cells were then incubated for one hour with 2µg/mL Hoechst solution (bisBenzimide H 33342 trihydrochloride, Sigma-Aldrich) and then washed three times with PBS. After, the images were collected with a 40X immersion objective in a confocal microscope on an inverted stand DMI6000 (Leica Microsystems, Mannheim, Germany). The excitation sources and detecting windows were 405nm (UV diode laser) / 410-520nm (PMT detector) for Hoechst; 543nm (Green Helium-Neon) / 548-652nm (PMT detector) for Cy3; 633nm (Red Helium-Neon) / 638-765nm (PMT detector) for Cy5; 543nm / 638-765 (PMT detector) for FRET. Images were then analyzed with Fiji software (version 1.54p). Background noise was subtracted and the different FRET efficiencies were calculated with the Fret and Colocalization Analyzer according to its user’s guide, on camera full-frame images. After analysis, the number of FRET-positive pixels was obtained for each image. The number of cells was obtained by counting the nuclei. For statistical analysis, the amount of FRET-positive pixels and FRET-negative pixels per image (total image size of 1024×1024 pixels) were calculated for the UNZIP and ZIP conditions and normalized for 100 cells. Then, a Chi-square with Yates’ correction was performed using these parameters between the 2 conditions.

### Statistical analysis and reproducibility

The experimental data presented are from the analysis of several independent experiments. No statistical method was used to predetermine sample size. No data were excluded from the analyses.

The experiments were not randomized. Statistical significance was inferred when necessary and formatted using GraphPad Prism 10.3.1 software. To confirm significance, p-values were calculated when analyzing at least three independent experiments. Exact distinct and independent experiments size is indicated in each legend (n). Results are presented as mean ± SD. To compare two groups, the parametric t test (two-sided) or the nonparametric Mann–Whitney test (two-sided) were used when residual distribution was Gaussian/normal or not respectively. If the comparison was done to a condition normalized to 1, the nonparametric Kolmogorov-Smirnov test was used. To compare more than two groups, one-way ANOVA (two-sided) was used and Gaussian/normal residual distribution was checked. If Gaussian/normal residual distribution was not assumed or not validated, Kruskal-Wallis test was used. To compare several conditions in two groups or more, 2way ANOVA was used and Gaussian/normal residual distribution was checked. If not indicated, the results are not significant. The details (number of independent replicates, statistical test used) are indicated in the legends of the figures.

## Data availability

All raw data and bioinformatic pipelines are available upon request to corresponding authors.

## Reporting summary

Further information on the research design is available in the Nature Portfolio Reporting Summary linked to this article.

## Acknowledgements

We thank Nathalie Dugot-Senant from the Histopathology platform and all the staff from Vect’UB, FACSility and One cell facilities at University of Bordeaux (TBMCore), Sandrine Hamon, Stephanie Lannelongue and all the BRIC administrative staff for administrative assistance and financial management, Bagatelle Hospital (Talence, France) for cord blood samples, Louis Domenach for his input on cystic fibrosis and Ray Cooke for copyediting the manuscript. C.T. is supported by funding from the AFM-Telethon (#24581). This work was supported by an AFM-Telethon grant (#24136), CoPOC ZIP-CRISPR (RVF23035GSA) and the French ANR grants GENOTOX (ANR-21-CE18-0002-01) and CRISPR SAFE (ANR-PRME-24-CE52).

## Author contributions

C.T., J.R., F.M.G. and A.B.: design and analysis of data; C.T., F.M.G. and A.B.: drafting of manuscript; A.B. and F.M.G.: supervision, funding acquisition, final approval of manuscript; C.T., J.R., M.R., V.M., S.F. and R.E.: performing editing experiments and associated analysis; V.M., C.T., C.B.: Nanopore sequencing bioinformatic analysis; C.T., I.L.G.: performing cell cycle analysis; I.L.G.: isolation of HSPCs; R.E. design and analysis of FRET experiment; M.L.: spectrofluorometry analysis; T.T., S.B.: obtention of pulmonary epithelial basal cells, nasal epithelial cells and bronchial epithelial cells; C.B., J.B,: helpful discussion in the field of CRISPR-Cas9; S.D., C.B., J.B.: proofreading of manuscript.

