## Supplementary file, figures and tables for "Improving efficiency of homology-directed repair with ZIP CRISPR"

Suppl Fig. S1

a

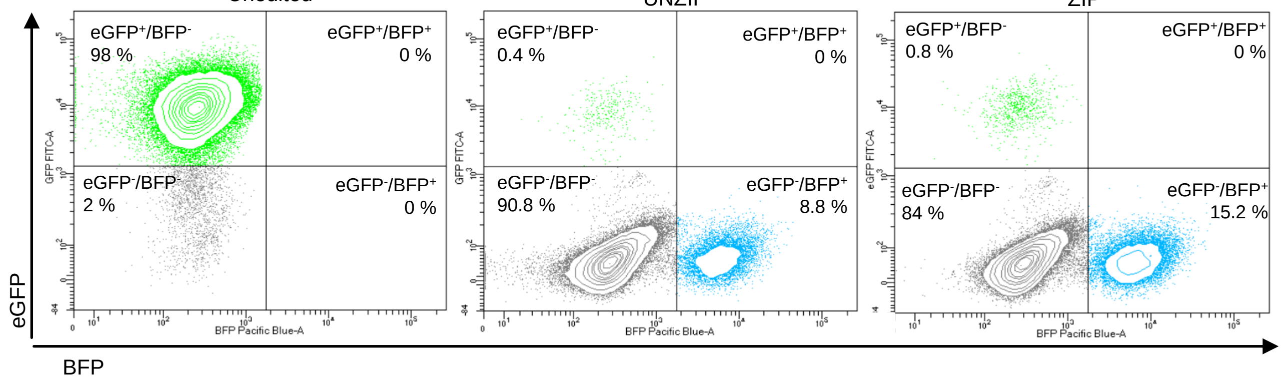


b

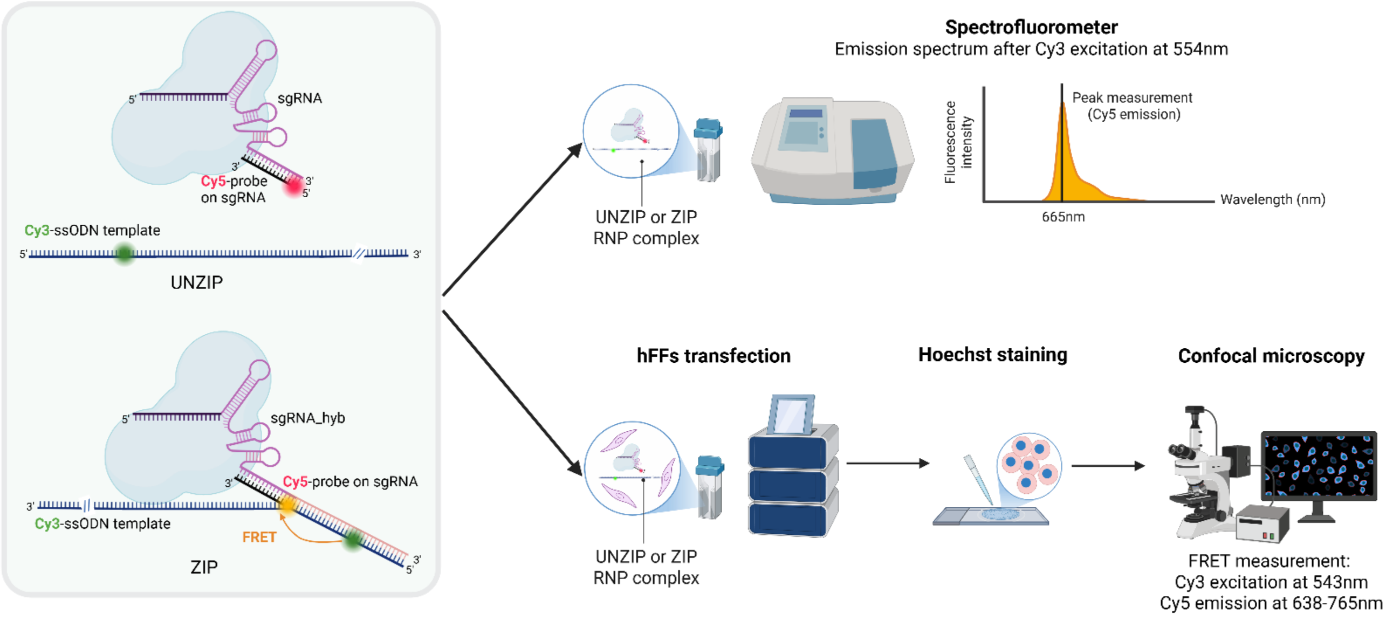


c

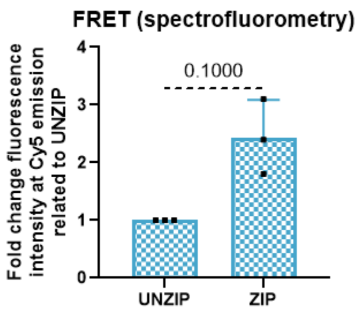


d

| | Usual dose | | | | Low dose | | | |
|---|---|---|---|---|---|---|---|---|
| | Cy3-ssODN | Cy5-probe on gRNA | UNZIP | ZIP | Cy3-ssODN | Cy5-probe on gRNA | UNZIP | ZIP |
| Cy3 | 29675 | 5585 | 52045 | 190539 | 11400 | 338 | 4194 | 16294 |
| Cy5 | 7331 | 244492 | 78755 | 619517 | 3213 | 18206 | 5882 | 13272 |

e

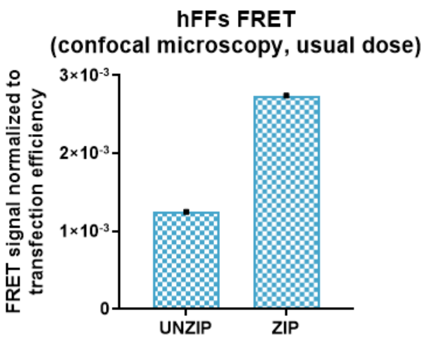

**Supplementary Fig. S1: ZIP CRISPR validation. a** Illustrative cytometry results of eGFP$^{+}$-HEK293T cells, unedited, edited by UNZIP or edited by ZIP**. b** Design of UNZIP and ZIP RNPs for FRET analysis and result analysis workflow. **c** Quantification of FRET signals observed by spectrofluorometry with UNZIP or ZIP systems (n=3). The FRET signal was measured by fluorescence intensity at 665 nm (Cy5 emission) after excitation at 554 nm (Cy3 excitation). Fold change indicates fold increase between UNZIP and ZIP conditions. The mean ± SD is shown. Statistical significance determined by Kolmogorov-Smirnov test. **d** Quantification of Cy3 or Cy5 signal observed by confocal microscopy in hFFs transfected with Cy3-ssODN, Cy5-probe on gRNA, UNZIP or ZIP systems at usual or low dose (n=1). The numbers of Cy3 or Cy5 positive-pixels normalized for 100 cells are indicated. **e** Quantification of FRET signal observed by confocal microscopy in hFFs transfected with ZIP or UNZIP systems at usual dose normalized on transfection efficiency (n=1). The ratio between the number of FRET-positive pixels normalized for 100 cells and the minimum between Cy3- and Cy5-positive pixels for 100 cells for each condition is represented. Schemas created with BioRender.com.

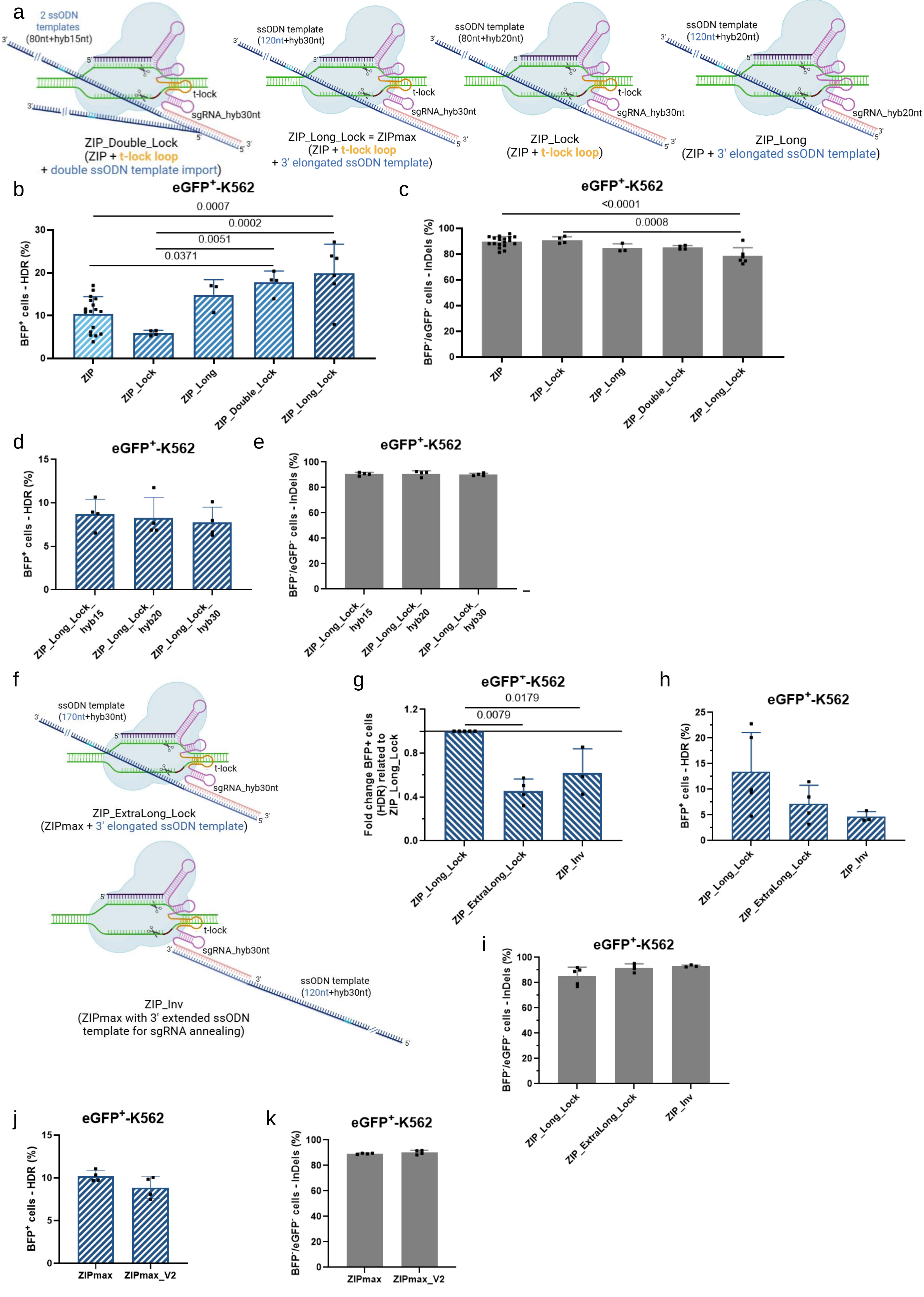
a
2 ssODN templates (80nt+hyb15nt)
t-lock
sgRNA_hyb30nt
ZIP_Double_Lock
(ZIP + t-lock loop
+ double ssODN template import)
ssODN template (120nt+hyb30nt)
ZIP_Long_Lock = ZIPmax
(ZIP + t-lock loop
+ 3' elongated ssODN template)
ssODN template (80nt+hyb20nt)
ZIP_Lock
(ZIP + t-lock loop)
ssODN template (120nt+hyb20nt)
sgRNA_hyb20nt
ZIP_Long
(ZIP + 3' elongated ssODN template)
b
eGFP+-K562
BFP+ cells - HDR (%)
0.0007
0.0002
0.0051
0.0371
ZIP
ZIP_Lock
ZIP_Long
ZIP_Double_Lock
ZIP_Long_Lock
c
BFP-/eGFP- cells - InDels (%)
<0.0001
0.0008
d
ZIP_Long_Lock_hyb15
ZIP_Long_Lock_hyb20
ZIP_Long_Lock_hyb30
e
f
ssODN template (170nt+hyb30nt)
ZIP_ExtraLong_Lock
(ZIPmax + 3' elongated ssODN template)
ssODN template (120nt+hyb30nt)
ZIP_Inv
(ZIPmax with 3' extended ssODN template for sgRNA annealing)
g
Fold change BFP+ cells (HDR) related to ZIP_Long_Lock
0.0179
0.0079
ZIP_ExtraLong_Lock
ZIP_Inv
h
i
j
ZIPmax
ZIPmax_V2
k

**Supplementary Fig. S2: Zip Editing development. a** Representation of RNPs used in the ZIP-Double (ZIP with two ssODN templates), ZIP_Long_Lock=ZIPmax (ZIP with additive t-lock loop in the gRNA and 3-'elongated ssODN template), ZIP_Lock (ZIP with additive t-lock loop in the gRNA), and ZIP_Long (ZIP with 3-'elongated ssODN template) import systems. **b** Flow cytometry quantification of HDR edition with Cas9 nuclease with ZIP (n=17), ZIP_Lock (n=4), ZIP_Long (n=3), ZIP_Double_Lock (n=4) or ZIP_Long_Lock (n=6) in eGFP$^{+}$-K562 cells. The proportions of cells expressing BFP (HDR-edited) one week after transfection are reported. The mean ± SD is shown. Statistical significance determined by one-way ANOVA. **c** Flow cytometry quantification of imprecise editing with Cas9 nuclease with ZIP (n=17), ZIP_Lock (n=4), ZIP_Long (n=3), ZIP_Double_Lock (n=4) or ZIP-Long-Lock (n=6) in eGFP$^{+}$-K562 cells. The proportions of cells expressing neither BFP nor eGFP (imprecisely-edited) one week after transfection are reported. The mean ± SD is shown. Statistical significance determined by one-way ANOVA. **d** Flow cytometry quantification of HDR edition with Cas9 nuclease with ZIP_Long_Lock_hyb15, ZIP_Long_Lock_hyb20 or ZIP_Long_Lock_hyb30 in eGFP$^{+}$-K562 cells (n=4). The proportions of cells expressing BFP (HDR-edited) one week after transfection are reported. The mean ± SD is shown. **e** Flow cytometry quantification of imprecise editing with Cas9 nuclease with ZIP_Long_Lock_hyb15, ZIP_Long_Lock_hyb20 or ZIP_Long_Lock_hyb30 in eGFP$^{+}$-K562 cells (n=4). The proportions of cells expressing neither BFP nor eGFP (imprecisely-edited) one week after transfection are reported. The mean ± SD is shown. **f** Representation of RNPs used in the ZIP_ExtraLong_Lock (ZIP_Long_Lock with 3-'over-elongated ssODN template) and ZIP_Inv (ZIP_Long_Lock with ssODN 3'-hybridization to the gRNA, instead of 5') import systems. **g** Flow cytometry quantification of HDR edition with Cas9 nuclease with ZIP_Long_Lock (n=4), ZIP_ExtraLong_Lock (n=4) and ZIP_Inv (n=3) in eGFP$^{+}$-K562 cells. The proportions of cells expressing BFP (HDR-edited) one week after transfection are reported. Fold changes indicate fold increases between ZIP_ExtraLong_Lock or ZIP_Inv relative to ZIP_Long_Lock. The mean ± SD is shown. Statistical significance to ZIP_Long_Lock determined by Kolmogorov-Smirnov test. **h** Flow cytometry quantification of HDR edition with Cas9 nuclease with ZIP_Long_Lock (n=5), ZIP_ExtraLong_Lock (n=4) and ZIP_Inv (n=3) in eGFP$^{+}$-K562 cells. The proportions of cells expressing BFP (HDR-edited) one week after transfection are reported. The mean ± SD is shown. **i** Flow cytometry quantification of imprecise editing with Cas9 nuclease with ZIP_Long_Lock (n=5), ZIP_ExtraLong_Lock (n=4) and ZIP_Inv (n=3) in eGFP$^{+}$-K562 cells. The proportions of cells expressing neither BFP nor eGFP (imprecisely-edited) one week after transfection are reported. The mean ± SD is shown. **j** Flow cytometry quantification of HDR edition with Cas9 nuclease with ZIPmax or ZIPmax_V2 in eGFP$^{+}$-K562 cells (n=4). The proportions of cells expressing BFP (HDR-edited) one week after transfection are reported. The mean ± SD is shown. **k** Flow cytometry quantification of imprecise editing with Cas9 nuclease with ZIPmax or ZIPmax_V2 in eGFP$^{+}$-K562 cells (n=4). The proportions of cells expressing neither BFP nor eGFP (imprecisely-edited) one week after transfection are reported. The mean ± SD is shown. Schemas created with BioRender.com.

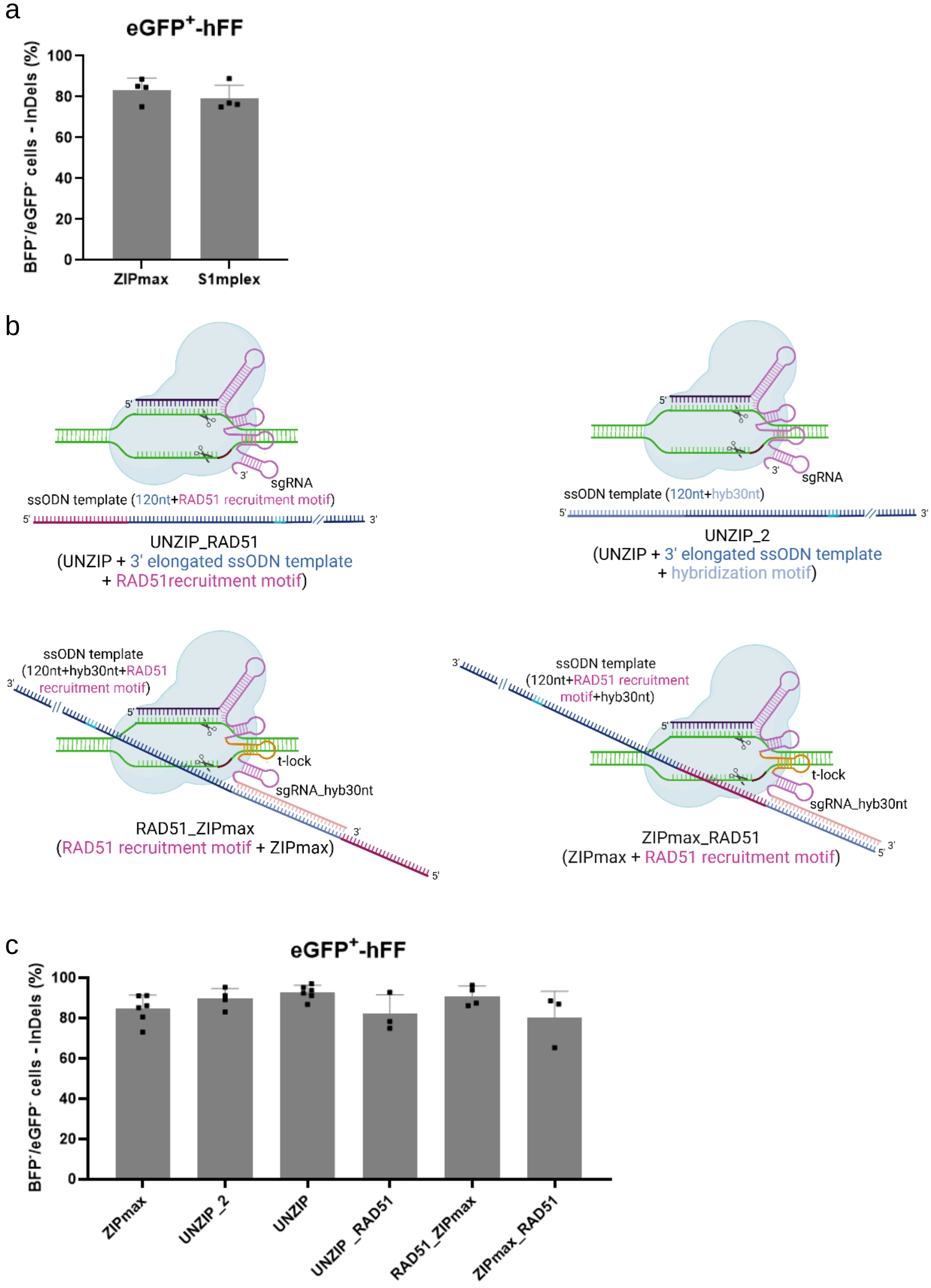


**Supplementary Fig. S3: ZIP CRISPR benchmarking. a** Flow cytometry quantification of imprecise editing with Cas9 nuclease with ZIPmax or S1mplex in eGFP$^+$-hFFs (n=4). The proportions of cells expressing neither BFP nor eGFP (imprecisely-edited) one week after transfection are reported. The mean ± SD is shown. **b** Representation of RNPs used in the UNZIP_RAD51, UNZIP_2, RAD51_ZIPmax and ZIPmax_RAD51 systems. **c** Flow cytometry quantification of imprecise editing with Cas9 nuclease with ZIPmax (n=4), UNZIP_2 (n=4), UNZIP (n=6), UNZIP_RAD51 (n=3), RAD51_ZIPmax (n=4) and ZIPmax_RAD51 (n=3) in eGFP$^+$-hFFs. The proportions of cells expressing neither BFP nor eGFP (imprecisely-edited) one week after transfection are reported. The mean ± SD is shown. Schemas created with BioRender.com.

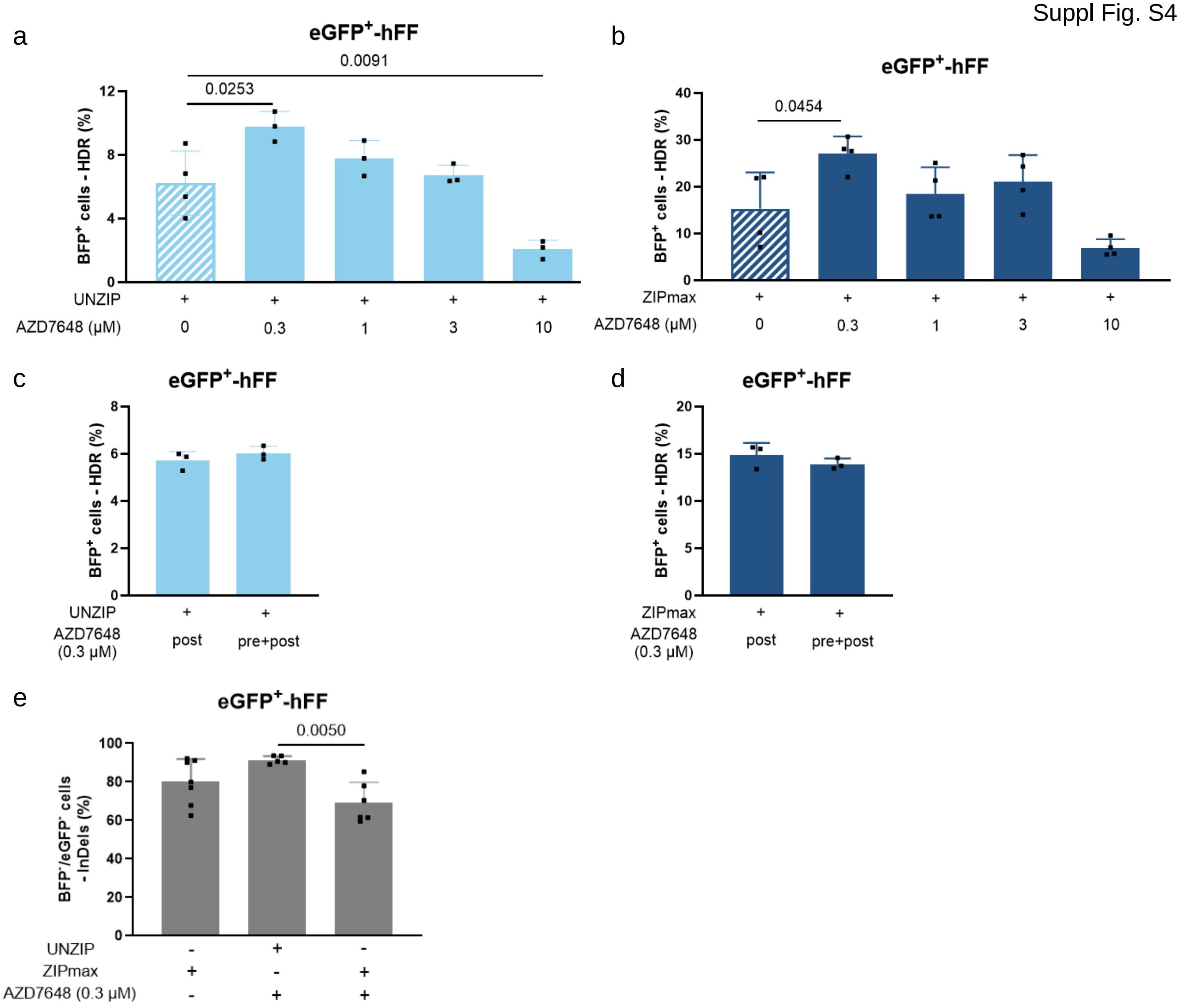


**Supplementary Fig. S4: Set up of AZD7648 use for comparison and combination to ZIP CRISPR. a and b** Flow cytometry quantification of the HDR edition with Cas9 nuclease with UNZIP (**a** n=3 or 4) or ZIPmax (**b** n=4) and with or without inhibition of NHEJ repair with different doses of AZD7648 in eGFP+-hFFs. The proportions of cells expressing BFP one week after transfection are reported. The mean ± SD is shown. Statistical significance determined by one-way ANOVA. **c and d** Flow cytometry quantification of the HDR edition with Cas9 nuclease with UNZIP (**c**) or ZIPmax (**d**) with inhibition of NHEJ repair with different timing of AZD7648 exposure in eGFP+-hFFs (n=3). The proportions of cells expressing BFP one week after transfection are reported. The mean ± SD is shown. Statistical significance determined by unpaired t-test. **e** Flow cytometry quantification of the imprecise edition with Cas9 nuclease with UNZIP (n=5) or ZIPmax (n=6 or 7) and with or without inhibition of NHEJ repair with AZD7648 in eGFP+-hFFs. The proportions of cells expressing neither BFP nor eGFP one week after transfection are reported. The mean ± SD is shown. Statistical significance determined by one-way ANOVA.

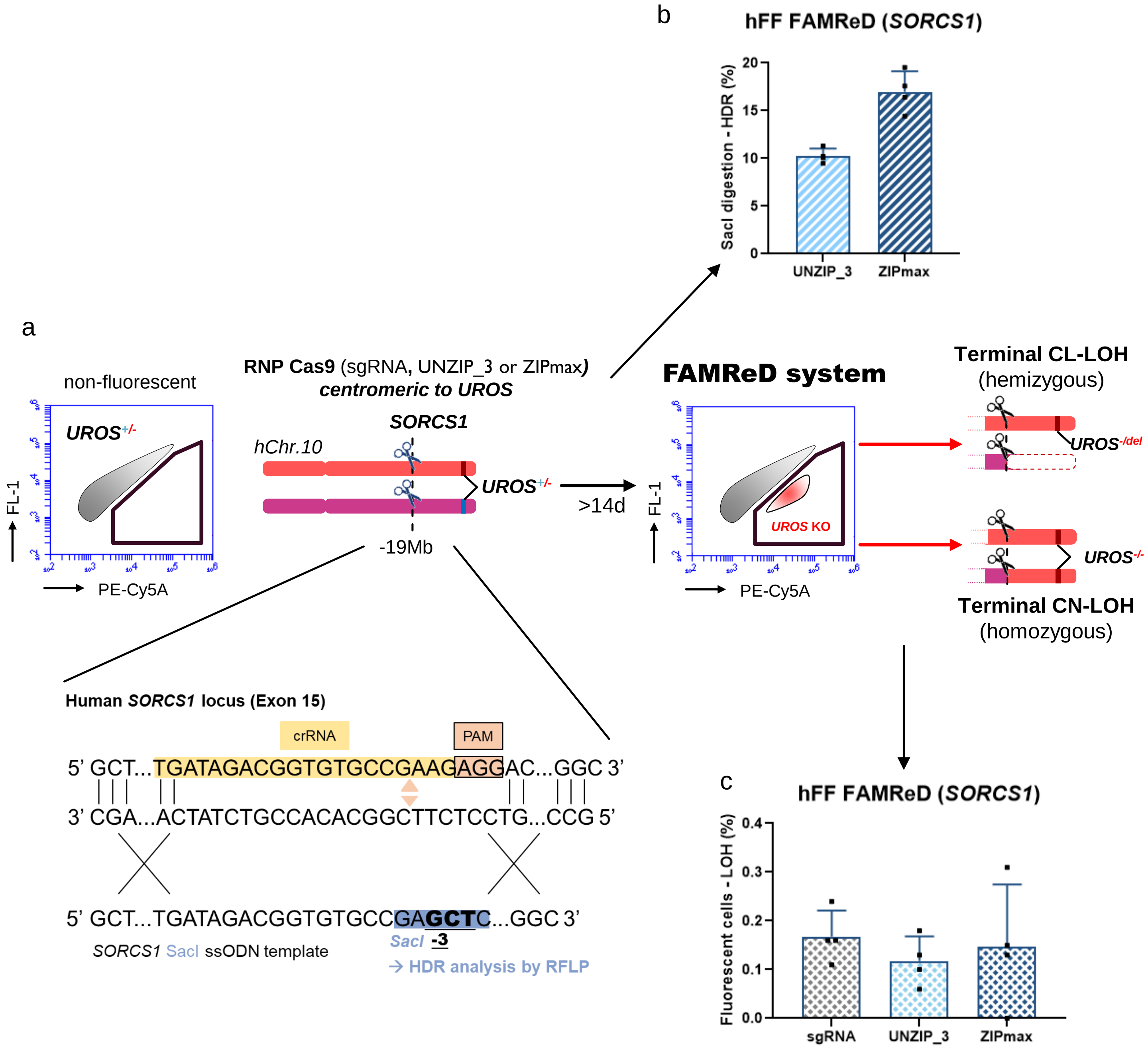


**Supplementary Fig. S5: FAMReD technology to detect loss of heterozygosity (LOH) after CRISPR editing. a** FAMReD system to detect and quantify LOH after CRISPR editing and editing strategy to edit *SORCS1* (insertion of *SacI* restriction site) in hFFs. **b** Capillary electrophoresis of *SacI* digestion products to quantify HDR editing with UNZIP_3 or ZIPmax (n=4). The mean ± SD is shown. **c** LOH quantification after CRISPR editing with sgRNA only, UNZIP_3 or ZIPmax (n=4). The mean ± SD is shown.

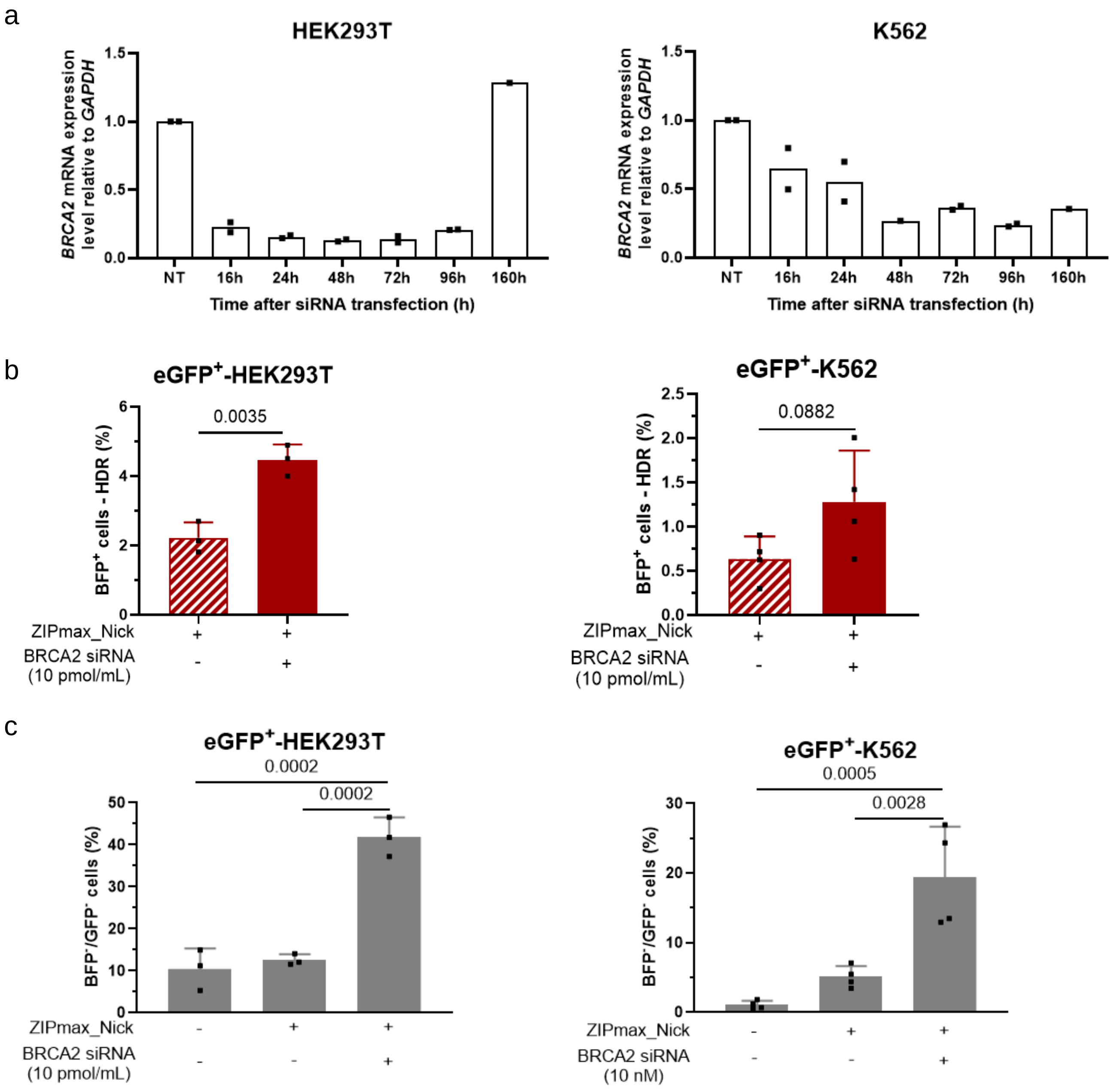


**Supplementary Fig. S6: Modulation of DNA repair pathways can be associated with ZIPmax_Nick to increase HDR editing. a** RTqPCR quantification of *BRCA2* mRNA expression levels over time after transfection of siRNA directed against *BRCA2* in HEK293T cells (left panel, n=2) or K562 cells (right panel, n=2). Values are normalized to *GAPDH* expression levels. The mean is shown. **b** Flow cytometry quantification of HDR edition with Cas9 nickase (ZIPmax_Nick) with or without inhibition of *BRCA2* by siRNA during edition in eGFP$^{+}$-HEK293T cells (left panel, n=3) or eGFP$^{+}$-K562 cells (right panel, n=4). The proportions of cells expressing BFP (HDR-edited) one week after transfection are reported. The mean ± SD is shown. Statistical significance determined by unpaired t-test. **c** Flow cytometry quantification of imprecise edition with Cas9 nickase (ZIPmax_Nick) with or without inhibition of *BRCA2* by siRNA in eGFP$^{+}$-HEK293T cells (left panel, n=3) or eGFP$^{+}$-K562 cells (right panel, n=4). The proportions of cells expressing neither BFP nor eGFP one week after transfection are reported. The mean ± SD is shown. Statistical significance determined by one-way ANOVA.

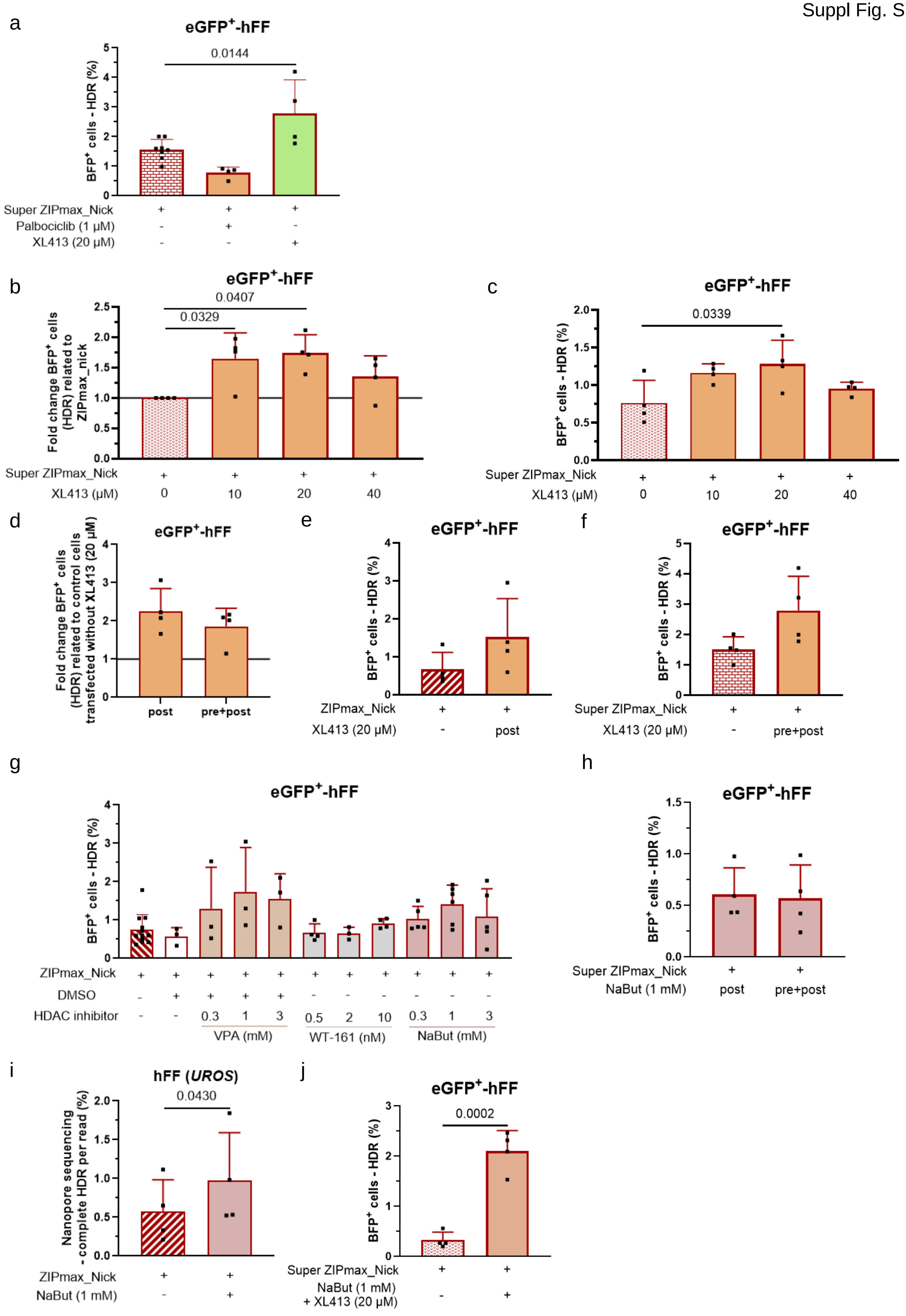
a
eGFP+-hFF
0.0144
BFP+ cells - HDR (%)
Super ZIPmax_Nick
Palbociclib (1 µM)
XL413 (20 µM)
b
eGFP+-hFF
0.0407
0.0329
Fold change BFP+ cells (HDR) related to ZIPmax_nick
Super ZIPmax_Nick
XL413 (µM)
0
10
20
40
c
eGFP+-hFF
0.0339
BFP+ cells - HDR (%)
Super ZIPmax_Nick
XL413 (µM)
d
eGFP+-hFF
Fold change BFP+ cells (HDR) related to control cells transfected without XL413 (20 µM)
post
pre+post
e
eGFP+-hFF
BFP+ cells - HDR (%)
ZIPmax_Nick
XL413 (20 µM)
post
f
eGFP+-hFF
BFP+ cells - HDR (%)
Super ZIPmax_Nick
XL413 (20 µM)
pre+post
g
eGFP+-hFF
BFP+ cells - HDR (%)
ZIPmax_Nick
DMSO
HDAC inhibitor
0.3
1
3
0.5
2
10
VPA (mM)
WT-161 (nM)
NaBut (mM)
h
eGFP+-hFF
BFP+ cells - HDR (%)
Super ZIPmax_Nick
NaBut (1 mM)
post
pre+post
i
hFF (UROS)
0.0430
Nanopore sequencing - complete HDR per read (%)
ZIPmax_Nick
NaBut (1 mM)
j
eGFP+-hFF
0.0002
BFP+ cells - HDR (%)
Super ZIPmax_Nick
NaBut (1 mM)
+ XL413 (20 µM)

**Supplementary Fig. S7: Set up of drug use for combination to ZIP CRISPR with Nickase. a** Flow cytometry quantification of HDR edition with Cas9 nickase (Super ZIPmax_Nick) with or without synchronization of eGFP$^+$-hFFs in G0/G1 phase with palbociclib or in S/G2/M phases with XL413 (n=4). The proportions of cells expressing BFP (HDR-edited) one week after transfection are reported. The mean ± SD is shown. Statistical significance determined by one-way ANOVA. **b** Flow cytometry quantification of HDR edition with nickase Cas9 (Super ZIPmax_Nick) with or without XL413 exposure at different doses (n=4). The proportion of cells expressing BFP (HDR-edited) one week after transfection is reported. Fold changes indicate fold increases between the conditions with and without drug. The mean ± SD is shown. Statistical significance determined by Kruskal-Wallis test. **c** Flow cytometry quantification of HDR edition with nickase Cas9 (Super ZIPmax_Nick) with or without XL413 exposure at different doses (n=4). The proportion of cells expressing BFP (HDR-edited) one week after transfection is reported. The mean ± SD is shown. Statistical significance determined by one-way ANOVA. **d** Flow cytometry quantification of HDR edition with nickase Cas9 with different timing of XL413 exposure (n=4). The proportion of cells expressing BFP (HDR-edited) one week after transfection is reported. Fold changes indicate fold increases between the conditions with and without drug. The mean ± SD is shown. **e and f** Flow cytometry quantification of HDR edition with nickase Cas9 (**e** ZIPmax_Nick or **f** Super ZIPmax_Nick) with or without XL413 and with different timing of XL413 exposure (n=4). The proportion of cells expressing BFP (HDR-edited) one week after transfection is reported. The mean ± SD is shown. **g** Flow cytometry quantification of HDR edition with nickase Cas9 (ZIPmax_Nick) with or without VPA (n=3), WT-161 (n=3) or NaBut (n=5) exposure at different doses. The proportion of cells expressing BFP (HDR-edited) one week after transfection is reported. **h** Flow cytometry quantification of HDR edition with nickase Cas9 (Super ZIPmax_Nick) with different timing of NaBut exposure (n=4). The proportion of cells expressing BFP (HDR-edited) one week after transfection is reported. The mean ± SD is shown. **i** Nanopore sequencing to quantify *UROS* HDR editing per read with Cas9 nickase (ZIPmax_Nick) with or without NaBut in hFFs (n=4). The mean ± SD is shown. Statistical significance determined by paired t-test. **j** Flow cytometry quantification of HDR edition with Cas9 nickase (Super ZIPmax_Nick) with or without NaBut and XL413 exposure in eGFP$^+$-hFFs (n=4). The proportions of cells expressing BFP (HDR-edited) one week after transfection are reported. The mean ± SD is shown. Statistical significance determined by unpaired t-test.

**Supplementary Table 1: gRNA sequences for CRISPR-Cas9 editing.**

| sgRNA | Sequence (5' to 3') |
|---|---|
| sgRNA eGFP (UNZIP, UNZIP_RAD51, UNZIP_2, UNZIP_Nick) | CTCGTGACCACCCTGACCTAGTTTTAGAGCTAGAAATAGCAAGTTAAAATAAGGCTAGTCCGTTATCAACTTGAAAAAGTGGCACCGAGTCGGTGCTTTT |
| sgRNA eGFP (ZIP, ZIP_Long) | CTCGTGACCACCCTGACCTAGTTTTAGAGCTAGAAATAGCAAGTTAAAATAAGGCTAGTCCGTTATCAACTTGAAAAAGTGGCACCGAGTCGGTGCTTTTCTGCCATCAAAGCGTGCTCAGTCT |
| sgRNA eGFP (ZIP_Lock, ZIP_Double_Lock, ZIPmax, RAD51_ZIP_max, ZIPmax_RAD51, ZIP_ExtraLong_Lock, ZIP_Inv, ZIPmax_Nick, Super ZIPmax_Nick) | CTCGTGACCACCCTGACCTAGTTTTAGAGCTAGAAATAGCAAGTTAAAATAAGGCTAGTCCGTTATCAACTTGGACTTCGGTCCAAGTGGCACCGAGTCGGTGCTTTTCTGCCATCAAAGCGTGCTCAGTCTGCTGCC |
| sgRNA eGFP (S1mplex) | CTCGTGACCACCCTGACCTAGTTTAAGAGCTATGCTGCGAATACGAGATGCGGCCGCCGACCAGAATCATGCCAAGTGCGTAAGATAGTCGCGGGTCGGCGGCCGCATCTCGTATTCGCAGCATAGCAAGTTTAAATAAGGCTAGTCCGTTATCAACTTGAAAAAGTGGCACCGTGACGGTGCTTT |
| sgRNA eGFP (ZIP_Long_Lock_hyb15) | CTCGTGACCACCCTGACCTAGTTTTAGAGCTAGAAATAGCAAGTTAAAATAAGGCTAGTCCGTTATCAACTTGGACTTCGGTCCAAGTGGCACCGAGTCGGTGCTTTTCTGCCATCAAAGCGT |
| sgRNA eGFP (ZIP_Long_Lock_hyb20) | CTCGTGACCACCCTGACCTAGTTTTAGAGCTAGAAATAGCAAGTTAAAATAAGGCTAGTCCGTTATCAACTTGGACTTCGGTCCAAGTGGCACCGAGTCGGTGCTTTTCTGCCATCAAAGCGTGCTCA |
| sgRNA eGFP (ZIPmax_V2) | CTCGTGACCACCCTGACCTAGTTTTAGAGCTAGAAATAGCAAGTTAAAATAAGGCTAGTCCGTTATCAACTTGGACTTCGGTCCAAGTGGCACCGAGTCGGTGCTTTTACGCCGGGTTACAACGAGCCTCTTTCTCCG |
| sgRNA HBB (UNZIP_3, ZIPmax) | GTAACGGCAGACTTCTCCTCGTTTTAGAGCTAGAAATAGCAAGTTAAAATAAGGCTAGTCCGTTATCAACTTGGACTTCGGTCCAAGTGGCACCGAGTCGGTGCTTTTCTGCCATCAAAGCGTGCTCAGTCTGCTGCC |
| sgRNA UROS (UNZIP_3, ZIPmax, UNZIP_Nick_3, ZIPmax_Nick) | GGAAGCAGCAGAGTTATGTTGTTTTAGAGCTAGAAATAGCAAGTTAAAATAAGGCTAGTCCGTTATCAACTTGGACTTCGGTCCAAGTGGCACCGAGTCGGTGCTTTTCTGCCATCAAAGCGTGCTCAGTCTGCTGCC |
| sgRNA CFTR_G542X_modeling (UNZIP) | GACAATATAGTTCTTGGAGAGTTTTAGAGCTAGAAATAGCAAGTTAAAATAAGGCTAGTCCGTTATCAACTTGAAAAAGTGGCACCGAGTCGGTGCTTTT |
| sgRNA CFTR_G542X_modeling (ZIPmax) | GACAATATAGTTCTTGGAGAGTTTTAGAGCTAGAAATAGCAAGTTAAAATAAGGCTAGTCCGTTATCAACTTGGACTTCGGTCCAAGTGGCACCGAGTCGGTGCTTTTCTGCCATCAAAGCGTGCTCATTCTGCCATCAAAGCGTGCTCA |
| sgRNA CFTR_G542X_correction (UNZIP_3, ZIPmax) | GACAATATAGTTCTTTGAGAGTTTTAGAGCTAGAAATAGCAAGTTAAAATAAGGCTAGTCCGTTATCAACTTGGACTTCGGTCCAAGTGGCACCGAGTCGGTGCTTTTCTGCCATCAAAGCGTGCTCAGTCTGCTGCC |
| sgRNA SORCS1 (UNZIP_3, ZIPmax) | TGATAGACGGTGTGCCGAAGGTTTTAGAGCTAGAAATAGCAAGTTAAAATAAGGCTAGTCCGTTATCAACTTGGACTTCGGTCCAAGTGGCACCGAGTCGGTGCTTTTCTGCCATCAAAGCGTGCTCAGTCTGCTGCC |

**Supplementary Table 2: HDR ssODN template sequences for CRISPR-Cas9 HDR editing (*eGFP* target).**

| HDR template | Sequence (5' to 3') |
|---|---|
| BFP (UNZIP, UNZIP_Nick) | AAGCTGCCCGTGCCCTGGCCCACCCTCGTGACCACCCTGAGCCACGGCGTGCAGTGCTTCGCCCGCTACCCCGACCACATGA |
| BFP (ZIP, ZIP_Lock, ZIPmax_hyb20) | TGAGCACGCTTTGATGGCAGAAGCTGCCCGTGCCCTGGCCCACCCTCGTGACCACCCTGAGCCACGGCGTGCAGTGCTTCGCCCGCTACCCCGACCACAT |
| BFP (ZIP_Double_Lock 1/2) | GGCAGCAGACTGAGCAAGCTGCCCGTGCCCTGGCCCACCCTCGTGACCACCCTGAGCCACGGCGTGCAGTGCTTCGCCCGCTACCCCGACCACAT |
| BFP (ZIP_Double_Lock 2/2, ZIP_Long_Lock_hyb15) | ACGCTTTGATGGCAGAAGCTGCCCGTGCCCTGGCCCACCCTCGTGACCACCCTGAGCCACGGCGTGCAGTGCTTCGCCCGCTACCCCGACCACAT |
| BFP (ZIP_Long, ZIPmax, UNZIP_2, ZIPmax_Nick) | GGCAGCAGACTGAGCACGCTTTGATGGCAGAAGCTGCCCGTGCCCTGGCCCACCCTCGTGACCACCCTGAGCCACGGCGTGCAGTGCTTCGCCCGCTACCCCGACCACATGAAGCAGCACGACTTCTTCAAGTCCGCCATGCCCGAAGGC |
| BFP-biotin (S1mplex) | Biotin-AAGCTGCCCGTGCCCTGGCCCACCCTCGTGACCACCCTGAGCCACGGCGTGCAGTGCTTCGCCCGCTACCCCGACCACATGA |
| BFP (UNZIP_RAD51) | AACTCCCCCTCCATCATTCACTGTAAGCTGCCCGTGCCCTGGCCCACCCTCGTGACCACCCTGAGCCACGGCGTGCAGTGCTTCGCCCGCTACCCCGACCACATGAAGCAGCACGACTTCTTCAAGTCCGCCATGCCCGAAGGC |
| BFP (RAD51_ZIPmax) | AACTCCCCCTCCATCATTCACTGTGGCAGCAGACTGAGCACGCTTTGATGGCAGAAGCTGCCCGTGCCCTGGCCCACCCTCGTGACCACCCTGAGCCACGGCGTGCAGTGCTTCGCCCGCTACCCCGACCACATGAAGCAGCACGACTTCTTCAAGTCCGCCATGCCCGAAGGC |
| BFP (ZIPmax_RAD51) | GGCAGCAGACTGAGCACGCTTTGATGGCAGAACTCCCCCTCCATCATTCACTGTAAGCTGCCCGTGCCCTGGCCCACCCTCGTGACCACCCTGAGCCACGGCGTGCAGTGCTTCGCCCGCTACCCCGACCACATGAAGCAGCACGACTTCTTCAAGTCCGCCATGCCCGAAGGC |
| BFP (ZIP_ExtraLong_Lock) | TGAGCACGCTTTGATGGCAGAAGCTGCCCGTGCCCTGGCCCACCCTCGTGACCACCCTGAGCCACGGCGTGCAGTGCTTCGCCCGCTACCCCGACCACATGAAGCAGCACGACTTCTTCAAGTCCGCCATGCCCGAAGGCTACGTCCAGGAGCGCACCATCTTCTTCAAGGACGACGGCACCTACAAGAC |
| BFP (ZIP_Inv) | AAGCTGCCCGTGCCCTGGCCCACCCTCGTGACCACCCTGAGCCACGGCGTGCAGTGCTTCGCCCGCTACCCCGACCACATGAAGCAGCACGACTTCTTCAAGTCCGCCATGCCCGAAGGCTGAGCACGCTTTGATGGCAG |
| BFP (ZIPmax_V2) | CGGAGAAAGAGGCTCGTTGTAACCCGGCGTAAGCTGCCCGTGCCCTGGCCCACCCTCGTGACCACCCTGAGCCACGGCGTGCAGTGCTTCGCCCGCTACCCCGACCACATGAAGCAGCACGACTTCTTCAAGTCCGCCATGCCCGAAGGC |
| BFP (Super ZIPmax_Nick) | GGCAGCAGACTGAGCACGCTTTGATGGCAGCACCTACGGCAAGCTGACCCTGAAGTTCATCTGCACCACCGGCAAGCTGCCCGTGCCCTGGCCCACCCTCGTGACCACCCTGAGCCACGGCGTGCAGTGCTTCGCCCGCTACCCCGACCA |

**Supplementary Table 3: HDR ssODN template sequences for CRISPR-Cas9 HDR editing (other targets).**

| HDR template | Sequence (5' to 3') |
|---|---|
| HBB (UNZIP_3) | TGACACAACTGTGTTCACTAGCAACCTCAAACAGACACCATGGTGCATCTGACTCCCGAGGAAAAATCCGCAGTCACTGCCCTGTGGGGCAAGGTGAACGTGGATGAAGTTGGTGGTGAG |
| HBB (ZIPmax) | GGCAGCAGACTGAGCACGCTTTGATGGCAGTGACACAACTGTGTTCACTAGCAACCTCAAACAGACACCATGGTGCATCTGACTCCCGAGGAAAAATCCGCAGTCACTGCCCTGTGGGGCAAGGTGAACGTGGATGAAGTTGGTGGTGAG |
| UROS (UNZIP_3, UNZIP_Nick_3) | CTGAAGATTACGGGGGACTCATTTTTACCAGCCCCAGAGCAGTGGAAGCAGCAGAGCTCTGTTTAGAGCAAAACAATAAAACTGAAGGTGAGGGTGGGTCTGCTGTGGATTCCACTGGAC |
| UROS (ZIPmax, ZIPmax_Nick) | GGCAGCAGACTGAGCACGCTTTGATGGCAGCTGAAGATTACGGGGGACTCATTTTTACCAGCCCCAGAGCAGTGGAAGCAGCAGAGCTCTGTTTAGAGCAAAACAATAAAACTGAAGGTGAGGGTGGGTCTGCTGTGGATTCCACTGGAC |
| CFTR_G542X_ modeling (UNZIP) | ACATCTCCAAGTTTGCAGAGAAAGACAATATAGTTCTTTAGGAGCTCGGAATCACACTGAGTGGAGGTCAACGAGCAAGA |
| CFTR_G542X_ modeling (ZIPmax) | TGAGCACGCTTTGATGGCAGAATTTTCTATTTTTGGTAATAGGACATCTCCAAGTTTGCAGAGAAAGACAATATAGTTTAGGAGCTCGGAATCACACTGAGTGGAGGTCAACGAGCAAGAATTTCTTTAGCAAGGTGAAT |
| CFTR_G542X_ correction (UNZIP_3) | TTCTATTTTTGGTAATAGGACATCTCCAAGTTTGCAGAGAAAGATAACATCGTCCTCGGAGAGGGTGGAATCACACTGAGTGGAGGTCAACGAGCAAGAATTTCTTTAGCAAGGTGAATA |
| CFTR_G542X_ correction (ZIPmax) | GGCAGCAGACTGAGCACGCTTTGATGGCAGTTCTATTTTTGGTAATAGGACATCTCCAAGTTTGCAGAGAAAGATAACATCGTCCTCGGAGAGGGTGGAATCACACTGAGTGGAGGTCAACGAGCAAGAATTTCTTTAGCAAGGTGAATA |
| SORCS1 (UNZIP_3) | GCTCTGAATGGCAGCTGGTCAAAGTAGATTACAAGTCCATTTTTGATAGACGGTGTGCCGAGCTCTACAGACCTTGGCAGCTGCACAGCCAGGTAGGAGGAAAAGAACTGAGACTAAGGC |
| SORCS1 (ZIPmax) | GGCAGCAGACTGAGCACGCTTTGATGGCAGGCTCTGAATGGCAGCTGGTCAAAGTAGATTACAAGTCCATTTTTGATAGACGGTGTGCCGAGCTCTACAGACCTTGGCAGCTGCACAGCCAGGTAGGAGGAAAAGAACTGAGACTAAGGC |

**Supplementary Table 4: PCR primer sequences.**

| PCR primers | Sequence (5' to 3') |
|---|---|
| HBB_424pb_F | CTGATGGTATGGGGCCAAGAG |
| HBB_424pb_R | GTCTCCACATGCCCAGTTTCT |
| UROS_405pb_F (RFLP analysis) | TAGTTCCAGGCACATAGTAAGCAC |
| UROS_405pb_R (RFLP analysis) | TCCCAAGGCAGAGTCTGTGA |
| UROS_1kb_F (Nanopore sequencing) | CTCTAATCCCAGGCTGCGTC |
| UROS_1kb_R (Nanopore sequencing) | GCCTCCGCTCATCAGTGTAA |
| CFTR_415pb_F (RFLP analysis) | CAGCAATGTTGTTTTTGACCAACT |
| CFTR_415pb_R (RFLP analysis) | ACCCACTAGCCATAAAACCCC |
| CFTR_1kb_F (Nanopore sequencing) | AGGTCGTGAGAATGAGGTGC |
| CFTR_1kb_R (Nanopore sequencing) | TGGAGTGGCAGGGTCTATGA |
| SORCS1_484pb_F | TGAACGCCCCACAAATGCTC |
| SORCS1_484pb_R | TTGGATCTGAGTGCTGAACTGG |

**Supplementary Table 5: Commands for nCRISPResso2 analysis.**

```
GUIDE="GACAATATAGTTCTTTGAGA"
AMPSEQ_REF="cagcaatgttgtttttgaccaactaaataaattgcatttgaaataatggagatgcaatgttcaaaatt
tcaactgtggttaaagcaatagtgtgatatatgattacattagaaggaagatgtgcctttcaaattcagattgagcatac
taaaagtgactctctaattttctatttttggtaatagGACATCTCCAAGTTTGCAGAGAAAGACAATATAGTTCTTGGAG
AAGGTGGAATCACACTGAGTGGAGGTCAACGAGCAAGAATTTCTTTAGCAAGgtgaataactaattattggtctagcaag
catttgctgtaaatgtcattcatgtaaaaaaattacagacatttctctattgctttatattctgtttctggaattgaaaa
aatcctggggttttatggctagtgggt"
AMPSEQ_HTZ="cagcaatgttgtttttgaccaactaaataaattgcatttgaaataatggagatgcaatgttcaaaatt
tcaactgtggttaaagcaatagtgtgatatatgattacattagaaggaagatgtgcctttcaaattcagattgagcatac
taaaagtgactctctaattttctatttttggtaatagGACATCTCCAAGTTTGCAGAGAAAGACAATATAGTTCTTTGAG
AAGGTGGAATCACACTGAGTGGAGGTCAACGAGCAAGAATTTCTTTAGCAAGgtgaataactaattattggtctagcaag
catttgctgtaaatgtcattcatgtaaaaaaattacagacatttctctattgctttatattctgtttctggaattgaaaa
aatcctggggttttatggctagtgggt"
HDRSEQ="cagcaatgttgtttttgaccaactaaataaattgcatttgaaataatggagatgcaatgttcaaaatttcaa
ctgtggttaaagcaatagtgtgatatatgattacattagaaggaagatgtgcctttcaaattcagattgagcatactaaa
agtgactctctaattttctatttttggtaatagGACATCTCCAAGTTTGCAGAGAAAGATAACATCGTCCTCGGAGAGGG
TGGAATCACACTGAGTGGAGGTCAACGAGCAAGAATTTCTTTAGCAAGgtgaataactaattattggtctagcaagcatt
tgctgtaaatgtcattcatgtaaaaaaattacagacatttctctattgctttatattctgtttctggaattgaaaaaatc
ctggggttttatggctagtgggt"
CRISPResso \
    --fastq_r1 "$fq" \
    --amplicon_seq "$AMPSEQ_HTZ,$AMPSEQ_REF" \
    --amplicon_name "HTZ,REF" \
    --flexiguide_seq "$GUIDE" --flexiguide_homology 90 \
    --ignore_substitutions \
    --expected_hdr_amplicon_seq "$HDRSEQ" \
    --default_min_aln_score 60 \
    --min_bp_quality_or_N 13 \
    --plot_window_size 20 \
    --min_frequency_alleles_around_cut_to_plot 0.1 \
    --quantification_window_size 10 \
    --quantification_window_center 0 \
    --cleavage_offset -3 \
    --exclude_bp_from_left 100 \
    --exclude_bp_from_right 100 \
    --min_average_read_quality 10 \
    -p 20 \
    --name "$name"
```

**Supplementary Table 6: RTqPCR primer sequences.**

| RTqPCR primers | Sequence (5’ to 3’) |
|---|---|
| BRCA2_RTqPCR_F | CCAAGTGGTCCACCCCAAC |
| BRCA2_RTqPCR_R | GGCTGAGACAGGTGTGGAAA |
| GAPDH_RTqPCR_F | CTGCACCACCAACTGCTTAG |
| GAPDH_RTqPCR_R | AGGTCCACCACTGACACGTT |

**Supplementary Table 7: gRNA, probes and HDR ssODN template sequences for FRET analysis.**

| Cy5-sgRNA Probe | Sequence (5’ to 3’) |
|---|---|
| UNZIP / ZIP | C*ACCGACTCGGTGCC |

| sgRNA | Sequence (5’ to 3’) |
|---|---|
| UNZIP | CTCGTGACCACCCTGACCTAGTTTTAGAGCTAGAAATAGCAAGTTAAAATAAGGC TAGTCCGTTATCAACTTGAAAAAGTGGCACCGAGTCGGTGCTTTT |
| ZIP | CTCGTGACCACCCTGACCTAGTTTTAGAGCTAGAAATAGCAAGTTAAAATAAGGC TAGTCCGTTATCAACTTGGACTTCGGTCCAAGTGGCACCGAGTCGGTGCTTTTC TGCCATCAAAGCGTGCTCAGTCTGCTGCC |

| Cy3-HDR template | Sequence (5’ to 3’) |
|---|---|
| UNZIP / ZIP | GGCAGCAGACTGAGCACGCTT*TGATGGCAGAAGCTGCCCGTGCCCTGGCCCA CCCTCGTGACCACCCTGAGCCACGGCGTGCAGTGCTTCGCCCGCTACCCCGAC CACATGAAGCAGCACGACTTCTTCAAGTCCGCCATGCCCGAAGGC |
